# A Spatiotemporal Notch Interaction Map from Membrane to Nucleus

**DOI:** 10.1101/2022.12.21.521435

**Authors:** Alexandre P. Martin, Gary A. Bradshaw, Robyn J. Eisert, Emily D. Egan, Lena Tveriakhina, Julia M. Rogers, Andrew N. Dates, Gustavo Scanavachi, Jon C. Aster, Tom Kirchhausen, Marian Kalocsay, Stephen C. Blacklow

## Abstract

Notch signaling relies on ligand-induced proteolysis to liberate a nuclear effector that drives cell fate decisions. The location and timing of individual steps required for proteolysis and movement of Notch from membrane to nucleus, however, remain unclear. Here, we use proximity labeling with quantitative multiplexed mass spectrometry to monitor the microenvironment of endogenous Notch2 after ligand stimulation in the presence of a gamma secretase inhibitor and then as a function of time after inhibitor removal. Our studies show that gamma secretase cleavage of Notch2 occurs in an intracellular compartment and that formation of nuclear complexes and recruitment of chromatin-modifying enzymes occurs within 45 minutes of inhibitor washout. This work provides a spatiotemporal map of unprecedented detail tracking the itinerary of Notch from membrane to nucleus after activation and identifies molecular events in signal transmission that are potential targets for modulating Notch signaling activity.

## INTRODUCTION

Notch signaling is an essential and conserved mechanism of cell-cell communication that controls normal development and maintains adult tissue homeostasis in a wide range of tissues and organ systems (*1, 2*). Mutations in Notch signaling components result in several human developmental disorders such as Alagille syndrome, caused by loss of function mutations of either *NOTCH2* or *JAGGED1* (*3, 4*), spondylocostal dysostosis (*5*), and Hajdu-Cheney disease (*6*). In addition, Notch mutations and/or deregulated Notch signaling are frequently found in cancer (*7*). Activating mutations of *NOTCH1* occur in more than 50% of T cell acute lymphoblastic leukemias (T-ALL), and similar activating mutations have been found in triple-negative breast cancer, adenoid cystic carcinoma, and tumors derived from pericytes or smooth muscle (*7*). On the other hand, Notch acts as a tumor suppressor in cutaneous squamous cell carcinomas (*8*), highlighting the complexities in targeting Notch for cancer therapy.

Notch proteins (Notch1-4 in mammals) are transmembrane receptors that transmit signals in response to canonical Delta-like or Jagged ligands (Jagged1, Jagged2, DLL1, DLL4) present on a signal-sending cell. Ligand binding initiates signal transduction by triggering a series of proteolytic cleavages of Notch in the receiver cell. The first ligand-induced cleavage is catalyzed by ADAM10 at a site called S2, external to the plasma membrane. S2-cleaved Notch molecules are then cleaved by gamma secretase at site S3, resulting in the release of the Notch intracellular domain (NICD). NICD subsequently translocates into the nucleus and forms a Notch transcription complex with the DNA-binding protein RBPJ and a MAML coactivator to induce the transcription of Notch target genes (see (*9*) and (*10*) for recent reviews).

While these fundamental steps required for Notch signaling have been defined, it is less certain where gamma secretase cleavage takes place in the cell, whether NICD moves from membrane to nucleus by active or passive transport, and how long it takes NICD to migrate from membrane to nucleus after gamma secretase cleavage. Real-time luciferase complementation assays using ectopic expression of Notch1 and RBPJ fusion proteins have shown that immobilized ligand stimulation results in nuclear complementation between 30 – 60 min after removal of a gamma secretase inhibitor (*11*), but analogous experiments have not been carried out at endogenous protein abundance. Previous reports have also reached different conclusions, for example, about whether gamma secretase cleavage occurs at the plasma membrane (*12, 13*) or in an intracellular compartment (*14*–*16*).

To address these questions, we mapped the microenvironment of NICD and characterized its interactions with effectors within a native cellular environment at endogenous expression levels after ligand stimulation in the presence of a gamma secretase inhibitor (GSI) and then as a function of time after inhibitor removal. We used CRISPR/Cas9 genome editing in SVG-A cells to fuse the Notch2 receptor to the engineered ascorbate peroxidase APEX2, which rapidly produces short-lived biotin-tyramide radicals that label proteins within a small radius (∼ 20 nm) and that can be rapidly quenched (*17*–*19*). We performed proximity labeling of the Notch2 microenvironment after ligand stimulation in the presence of a GSI and at different timepoints after inhibitor washout to identify changes in protein enrichment as a function of time by quantitative multiplexed proteomics. The dynamics of labeling enrichment of distinct plasma membrane, cytosolic, and nuclear proteins define the microenvironment of Notch2-APEX2 during its passage from the plasma membrane to the nucleus. Our studies show that gamma secretase cleavage of Notch2 to produce NICD2 occurs in an intracellular compartment, that passage of NICD2 through the cytoplasm is associated with transient enrichment of membrane cortical and cytoskeletal proteins, and that formation of nuclear complexes and recruitment of chromatin-modifying enzymes occurs within 45 minutes of inhibitor washout. This work provides a spatiotemporal map of unprecedented detail tracking the itinerary of Notch from membrane to nucleus after metalloprotease cleavage and identifies events in signal transmission that are potential targets for modulating Notch activity.

## RESULTS

### System validation and genome engineering for Notch2 proximity labeling in SVG-A cells

To track the movement of Notch as a function of time in response to signal induction, we labeled proteins in the Notch microenvironment using a Notch2-APEX2 ascorbate peroxidase fusion protein (*20*–*22*). We investigated the response associated with the Notch2/Jagged1 (Jag1) receptor-ligand pair because this pairing transmits signals that are required for normal development (*1*), as highlighted by Alagille syndrome, a multiorgan disorder caused by loss-of-function mutations in either *NOTCH2* or *JAGGED1* (*3, 4*). To achieve the precise synchronization necessary for time-resolved proximity labeling, we used immobilized Jag1 as an activating ligand in concert with a potent, specific gamma secretase inhibitor, the effects of which can be rapidly reversed by simple washout (*23*). We selected SVG-A human fetal astrocytes as our receptor-expressing cells (“Notch” or “receiver” cells) because they express abundant Notch2 endogenously and express low amounts of the other Notch receptors (Fig. S1A). We confirmed that Notch2 is responsible for the Notch transcriptional response in these cells by knocking out Notch2 using CRISPR/Cas9 genome editing (*24*): the absence of Notch2 effectively abolishes Notch reporter gene activity in a signaling assay using immobilized Jag1 as ligand (Fig. S1B-C), confirming both that these cells are responsive to Jag1 and that the reporter signal in these cells is a consequence of Notch2 activation.

To ensure that our studies were performed at natural receptor abundance, we used CRISPR/Cas9 to add an APEX2-HA coding sequence to the 3’ end of *NOTCH2*, creating a fusion gene at the endogenous locus encoding Notch2 fused to APEX2-HA at its C-terminus (Fig. S1D). The cassette for homologous recombination also contained a T2A sequence followed by a sequence encoding the mNeonGreen fluorescent protein, allowing us to isolate single cells positive for the desired genomic insertion by FACS. We confirmed that the Notch2-APEX2-HA fusion protein matures similarly to wild-type Notch2 in parental cells (Fig. S1E), is expressed at similar levels on the cell surface (Fig. S1F-G), retains signaling activity in response to immobilized Jag1 comparable to wild-type Notch2 in a reporter gene assay, and is silenced similarly to wild-type Notch2 by inhibitors of ADAM10 and gamma secretase cleavage (Fig. S1H). Western blot analysis also confirmed that biotinylation of proteins across a wide range of molecular weights was only observed in cells carrying the APEX2 fusion protein (Fig. S1I).

### Time-resolved Notch2 proximity labeling with the APEX2 fusion protein

Because soluble ligands do not activate Notch, it is not possible to use acute addition of the ligand to initiate the signal. To try to mimic an acute initiation event, we attempted to use removal of a selective ADAM10 inhibitor GI254023X (*25*) to serve as the event initiating S2 cleavage induced by immobilized ligand. If this approach were successful, it would have also allowed us to track dynamic events beginning with the S2 cleavage step generating the NEXT fragment. Unfortunately, however, this approach suffered from two complications that precluded the acquisition of time resolved proteomic information about signaling. First, “leaky” Notch2 proteolysis still occurred in the presence of the inhibitor, likely because ADAM17 or another protease can substitute for ADAM10 in catalyzing S2 cleavage to generate the NEXT fragment and second, the dissociation rate of GI254023X from ADAM10 was slow, and as a result the synchronization of events was poorer when this inhibitor was used (Fig. S2A). It was necessary, therefore, to use removal of a GSI to achieve the temporal resolution required for dynamic analysis of the Notch molecular neighborhood.

We thus cultured Notch2-APEX2-HA knock-in cells on culture plates containing immobilized Jag1 (*26*–*28*) overnight (16 h) in the presence of the gamma secretase inhibitor Compound E (GSI) and analyzed protein biotinylation as a function of time after GSI removal by mass spectrometry (Fig. 1A-B). We used a condition without ligand to serve as a baseline to define the molecular neighborhood of Notch2 in the absence of ligand. For the other conditions, we incubated the cells on immobilized ligand in the presence of GSI, set the time of inhibitor removal (when media containing GSI was replaced with GSI-free media) to t=0, and followed the time course of the Notch2 molecular neighborhood as a function of time after GSI washout (Fig. 1B). The cells were never treated with trypsin or EDTA, nor were they ever detached from the plate or resuspended, because all of these manipulations can activate Notch. We confirmed that GSI washout resulted in accumulation of NICD2 (*i.e*. S3-processed Notch2) over a two-hour time course (Fig. S2B), performed proximity labeling using biotin phenol and hydrogen peroxide at various time points up to 2 h, and by Western blot observed specific labeling of cohorts of biotin-labeled proteins that changed with time (Fig. S2C).

**Fig. 1.**
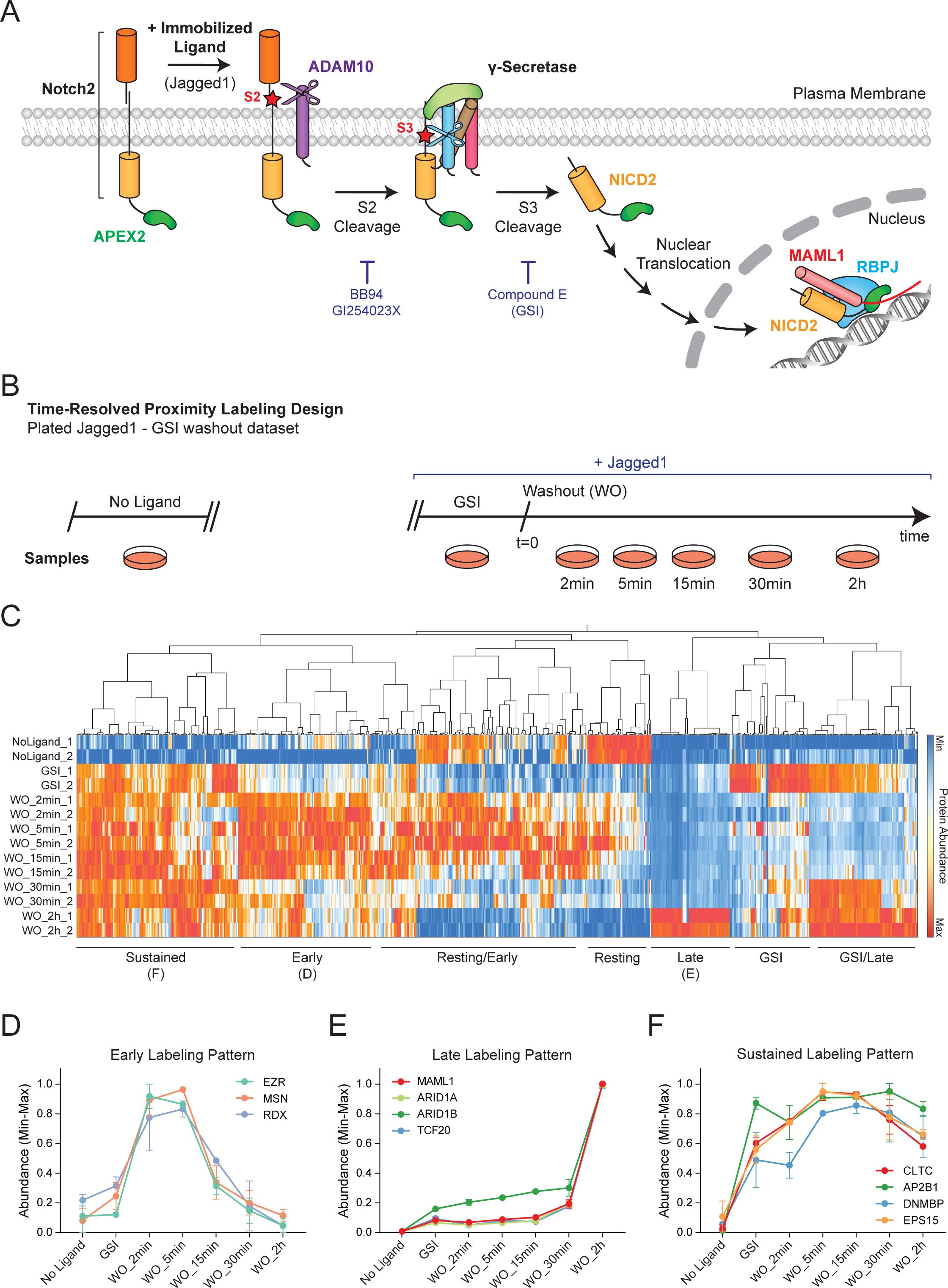
Design, experimental procedure, and overall kinetic profiles of time-resolved Notch2-APEX2 proximity labeling in SVG-A cells. (**A**) Schematic of key steps in Notch2 signaling induced by immobilized Jag1 ligand. Jag1 stimulates Notch2 proteolysis at S2 by ADAM10, followed by gamma secretase cleavage at S3. The S3-cleaved Notch2 intracellular domain (NICD2) transits to the nucleus and associates with RBPJ and the transcriptional co-activator MAML1 to induce the expression of target genes. BB94 and GI254023X (also referred as GI25X) are inhibitors of ADAM10, and Compound E (referred to as GSI) is a potent inhibitor of gamma secretase. (**B**) Schematic showing the design for time-resolved proximity labeling by Notch2-APEX2 using plated Jag1 as ligand and washout of GSI at time t=0. (**C**) Heatmap of hierarchical clustering of Notch2-APEX2 proximity labeling as a function of time after washout. Clustering of the relative abundance of each identified protein (columns) as a function of time (rows) was performed using Ward’s minimum variance method. Color palette representing the relative abundance for each protein (minimum to maximum) is shown on the right. (**D-F**) Kinetic profiles of representative proteins showing an early (**D**), late (**E**), or sustained (**F**) labeling pattern.

Mass spectrometry (MS) analysis of biotinylated proteins labeled by Notch2-APEX2 (Fig. S3A) determined the temporal profiles of labeling for 980 proteins, which displayed different dynamic labeling patterns after Notch activation. The high correlation between replicates (Fig. S3B) and the clustering of replicates in principal component analysis (Fig. S3C) attests to the reproducibility of our experimental system. When referenced to the t=0 timepoint, the transcriptional coactivator MAML1, an essential component of the Notch transcriptional complex (*29, 30*), was not significantly enriched at early timepoints after GSI washout (<30 min) but became the most significantly enriched protein two hours after GSI washout (see below), a finding independently verified by Western blot (Fig. S3D). This observation confirmed nuclear translocation of NICD2 after GSI removal and highlighted the dynamic nature of the NICD2 microenvironment as a function of time. We also note that our MS workflow did not identify all known Notch associated proteins in this study, as MS proteomics often does not capture all protein analytes present. For example, RBPJ, the transcription factor bound by NICD in the transcriptional activation complex, was not detected, even though RBPJ was indeed biotinylated as early as 30 min after GSI washout with a peak of relative abundance at 2 h after washout, as judged by Western blot (Fig. S3E).

Hierarchical clustering of the relative abundance of each labeled protein as a function of time led to the identification of seven distinct patterns of enrichment based on Ward’s hierarchical clustering method (Fig. 1C). These seven clusters show maximum labeling: 1) in the absence of ligand, 2) with ligand in the presence of GSI, 3) early (2-5 min) in the washout time course (Fig. 1D), 4) 2 h after washout (Fig. 1E), 5) throughout the time course in the presence of ligand, but not when ligand is absent (Fig. 1F), 6) with ligand in the presence of GSI and at later time points, and 7) both in the absence of ligand and at early time points. Full quantification data for proteins identified in this dataset are found in Data File S1.

We also co-cultured JAG1-expressing (A673) cells with Notch2-expressing cells in the presence of GSI followed by inhibitor washout, but the presence of ligand-expressing cells led to increased noise in studies spanning two different time windows (Fig. S4A-C). The trends seen in the data sets from these efforts are fully consistent with the results of the GSI washout using immobilized ligand, but the signals are much noisier and incomplete (Fig. S5A-E). MAML1 was not detected at the 2h timepoint, for example, in the co-culture experiments. The use of immobilized ligand was the only approach that eliminated interference from ligand-expressing cells, a decision necessary to achieve sufficient signal-to-noise for a robust time-resolved analysis. Full quantification data for proteins identified in these datasets are found in Data Files S2 and S3.

### Notch2 is internalized after S2 but before S3 cleavage

Not surprisingly, the first cluster, which exhibits maximum labeling enrichment in the absence of immobilized ligand (*i.e*. when Notch is unstimulated; Fig. 1C), is characterized by proteins that reside at the plasma membrane (Fig. 2A-B), where Notch encounters its transmembrane ligands present on neighboring cells. This group of proteins includes EGFR (Epidermal growth factor receptor), the folate carrier SLC19A1, ERBIN (Erbin), GPRC5A (Retinoic acid-induced protein 3), DBN1 (Drebrin), and surface-associated proteins such as the catenins CTNNB1 and CTNND1.

**Fig. 2.**
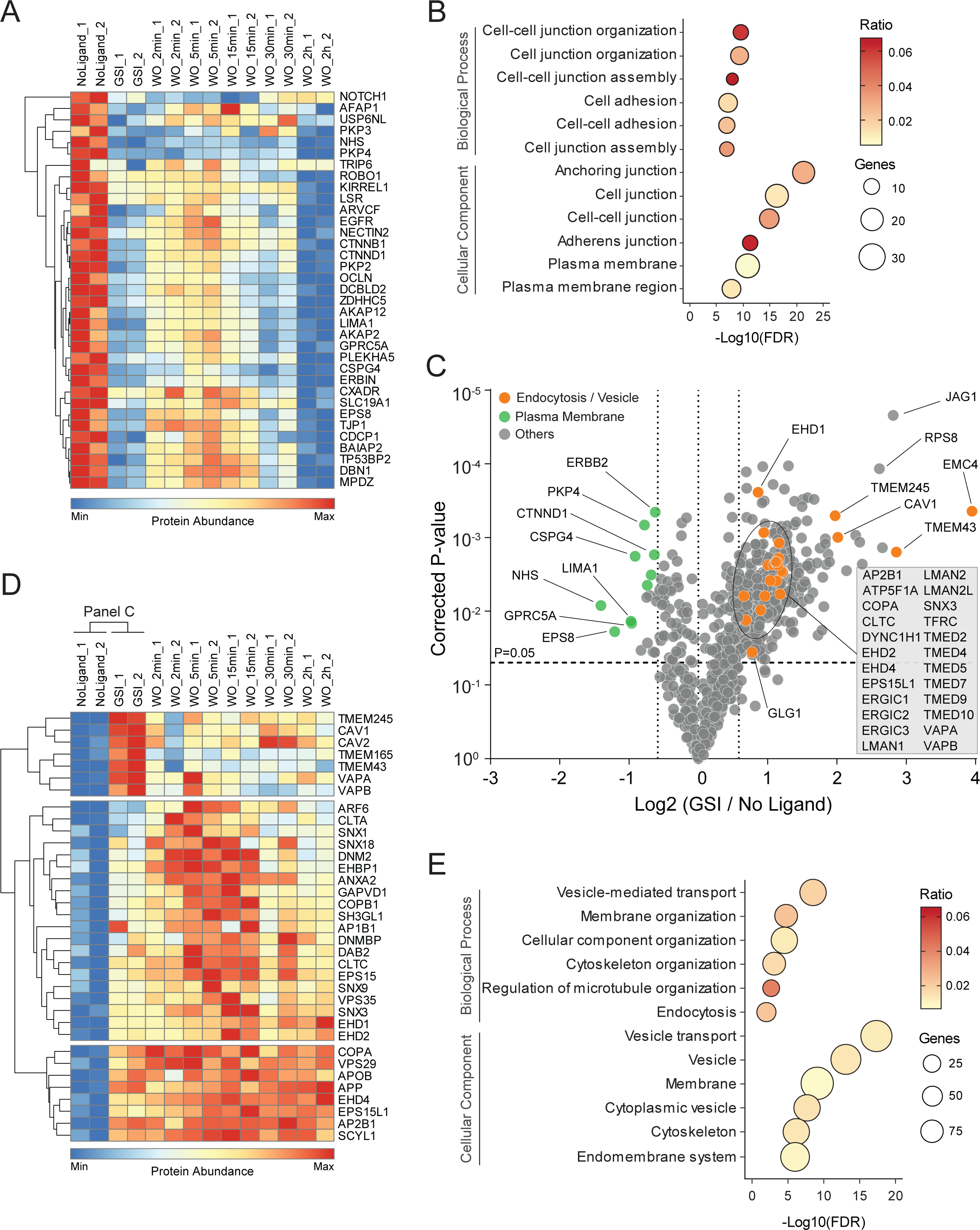
Changes in the Notch2 microenvironment upon stimulation by ligand in the presence of GSI. (**A**) Heatmap of hierarchical clustering of proteins characterized by peak relative abundance in conditions without Notch stimulation by ligand. (**B**) Gene Ontology terms for proteins significantly enriched in panel A. (**C**) Volcano plot comparing relative abundance of proteins enriched upon Jag1 stimulation in GSI compared to no ligand stimulation. Significantly enriched proteins (p-value ≤ 0.05, FC ≥ 1.5) related to endocytosis and/or vesicular-mediated transport are labeled in orange, whereas significantly downregulated proteins that localize to the plasma membrane are labeled in green. P-values are Benjamini-Hochberg corrected. (**D**) Heatmap focused on proteins related to endocytosis and vesicular transport identified in the time-resolved Notch2-APEX2 proximity labeling analysis. (**E**) Gene Ontology terms for proteins significantly enriched in panels C and D.

In the presence of GSI, ligand binding induces metalloprotease cleavage, but gamma secretase catalyzed cleavage at S3 is blocked, preventing liberation of NICD2 from the membrane. Remarkably, when GSI is present, plasma-membrane associated proteins, which are enriched in the first cluster when ligand is not present, become depleted when compared to unstimulated cells (Fig. 2C). Instead, proteins that exhibit maximal enrichment in their proximity to Notch2 after ligand exposure in the presence of GSI are predominantly associated with vesicular or endosomal compartments and the endocytic machinery (Fig. 2C-E). Core components participating in clathrin-mediated endocytosis (CME) including clathrin, AP-2, dynamin, and EPS15 are enriched (Fig. 1F and 2D) as is the transferrin receptor, which enters cells through CME, and other proteins implicated in vesicular trafficking such as EHD1, SEC22B, and SNX3 (Fig. 2C-E). In addition, there is a substantial enrichment of vesicle-associated TMED (transmembrane emp24 domain) proteins upon ligand stimulation (Fig. 2C-D), specifically TMEM165 and TMEM43. VAPA, and VAPB, two vesicle-associated proteins (*31, 32*), also show maximum enrichment in the presence of GSI (Fig. 2D). Another group of proteins showed labeling enrichment in the presence of GSI that persisted after washout (Fig. 2D). This group includes proteins related to endocytosis and vesicular trafficking (Fig. 2E), as well as the amyloid-beta precursor protein (APP; Fig. 2D), a well-known substrate of the gamma secretase complex (*33*). Finally, proteins related to endocytosis and components of the clathrin-mediated endocytic machinery (Fig. 2D) exhibited a significant increase in abundance upon ligand stimulation, with variable enrichment after GSI washout (Fig. 1F and 2D).

The comparison between the no ligand and GSI conditions suggests that Notch2 undergoes internalization after ligand-induced ADAM10 cleavage has occurred at site S2. To evaluate this possibility using a complementary approach, we monitored the subcellular localization of Notch2 by immunofluorescence after washout of GI254023X (GI25X), a potent ADAM10 inhibitor (*34*), in the absence or presence of the dynamin inhibitor hydroxy-dynasore, which blocks endocytosis (*35*–*37*). Cells were incubated overnight on immobilized Jagged1 in the presence of GI25X, and the inhibitor was then removed to allow S2 cleavage of ligand-bound Notch2 in the absence or presence of hydroxy-dynasore. As anticipated, washout of GI25X in the absence of the dynamin inhibitor resulted in nuclear accumulation of NICD2 by 2 h after washout (Fig. 3A-C), indicating that S2 cleavage, S3 cleavage, and nuclear translocation had occurred. However, washout of GI25X in the presence of hydroxy-dynasore (dynasore-OH) significantly impaired the nuclear accumulation of NICD2 (Fig. 3A-C). In addition, the inhibition of endosomal acidification by bafilomycin-A1 (BafA1), a specific vacuolar H^+^-ATPase inhibitor (*38*–*40*), or by chloroquine (*41*), also impaired Notch2 nuclear accumulation after GI25X washout (Fig. 3A-C), further suggesting that S3 cleavage required access to an acidified intracellular compartment. None of these inhibitors significantly modified the subcellular localization of unstimulated Notch2 (Fig. S6A), consistent with the conclusion that they act after ligand-induced S2 cleavage of Notch2.

**Fig. 3.**
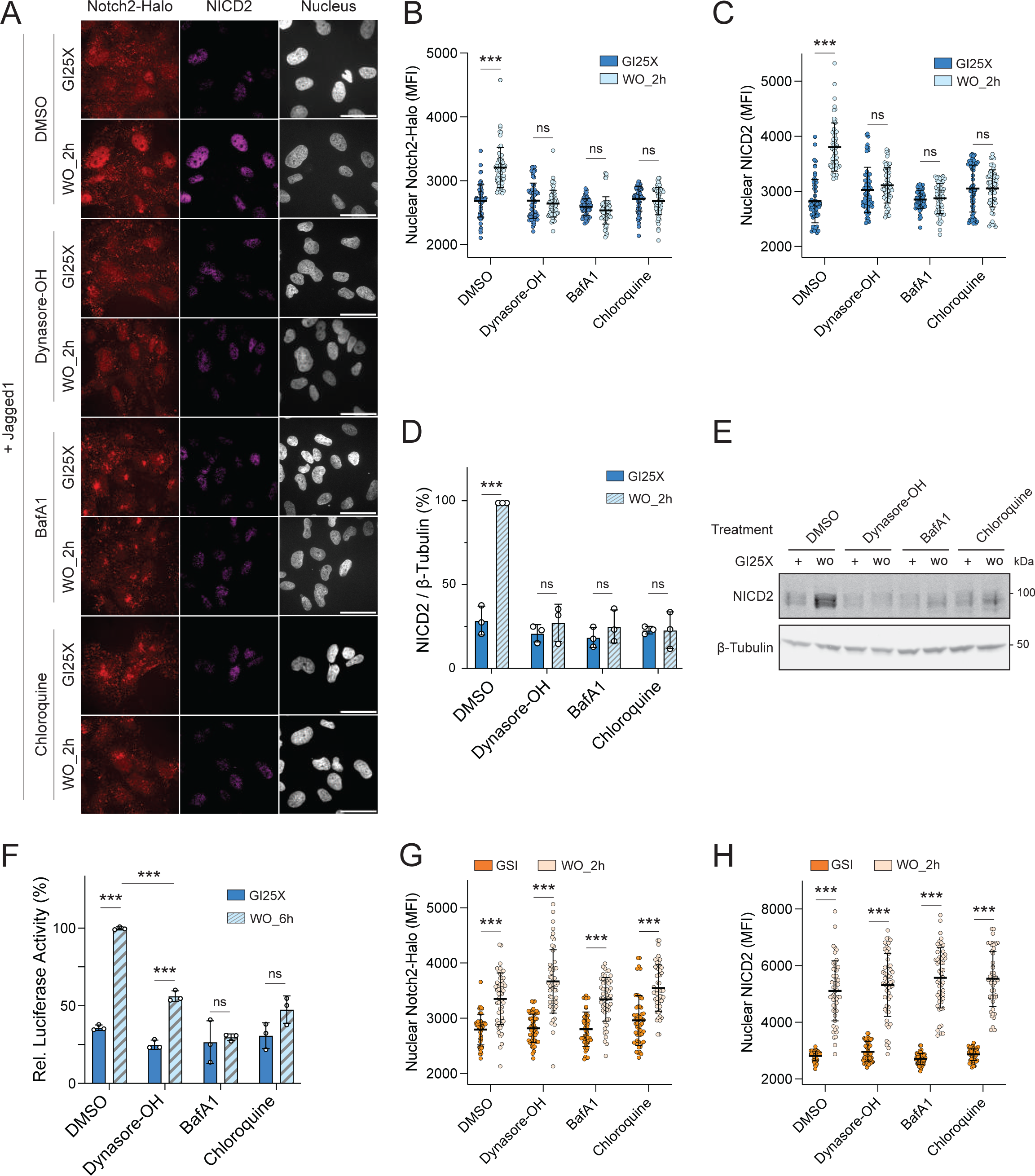
Effects of acidification and endocytosis inhibitors on Notch2 activity after removal of ADAM10 or gamma secretase inhibitors. (**A**) Representative images of Jag1-stimulated SVG-A Notch2-HaloTag cells showing the cellular distribution of Notch2-HaloTag (Notch2-Halo) and (S3-cleaved) NICD2 after removal of the ADAM10 inhibitor GI254023X in the absence or presence of hydroxy-dynasore (20 µM), bafilomycinA1 (BafA1; 25 nM), or chloroquine (50 µM). The HaloTag was labeled with JaneliaFluorX549 HaloTag ligand and NICD2 was stained with an anti-NICD2 primary antibody and anti-rabbit secondary antibody conjugated to Alexa Fluor 647. Nuclei were identified by DAPI staining. Scale bars: 20 µm. (**B-C**) Quantification of mean fluorescence signal intensity (MFI) in the nucleus for Notch2-Halo (**B**) and for NICD2 (**C**) for the imaging data presented in panel A for a total of 90 cells. (**D**) Quantification of Western blot data for NICD2 abundance after GI254023X washout in the presence of hydroxy-dynasore (20 µM), bafilomycinA1 (BafA1; 25 nM), or chloroquine (50 µM). (**E**) Representative Western blot for NICD2 quantified in panel D (see Fig. S7 for NICD1 and NICD2 generation in other cell lines). (**F**) Notch luciferase reporter assay. Parental SVG-A cells were stimulated by immobilized Jag1 overnight in the presence of GI254023X, and the relative luciferase activity was measured 6 h after removal of the ADAM10 inhibitor GI254023X in the absence or presence of hydroxy-dynasore (20 µM), bafilomycinA1 (BafA1; 25 nM), or chloroquine (50 µM). (**G-H**) Quantification of mean fluorescence signal intensity (MFI) in the nucleus (see fig. S6B for imaging data) for Notch2-Halo (**G**) and activated NICD2 (**H**) after removal of the GSI compound E (100 nM) in the absence or presence of hydroxy-dynasore (20 µM), bafilomycinA1 (BafA1; 25 nM), or chloroquine (50 µM) for a total of 90 cells. All data presented in this figure are from three independent experiments (n=3) and are presented as mean ± SD. *p ≤ 0.05, *p ≤ 0.01 and ***p ≤ 0.001. Two-way ANOVA followed by Tukey’s pairwise comparison was used for statistical comparisons.

The generation of the S3-cleaved form of Notch2 (NICD2) after GI25X washout was also evaluated by Western blotting, using an antibody that specifically recognizes the N-terminal epitope of NICD2 (*42*). As expected, washout of GI25X resulted in an increase in NICD2 abundance that was also greatly impaired by inhibition of endocytosis or vesicular acidification (Fig. 3D-E). Finally, the effect of inhibiting endocytosis or vesicular acidification on Notch transcriptional activity was investigated using a well-characterized luciferase reporter assay (*29, 43, 44*). Washout of GI25X induced a Notch-dependent transcriptional response that was substantially reduced by the presence of hydroxy-dynasore, BafA1, or chloroquine (Fig. 3F). In contrast, however, when GSI was washed out, NICD2 accumulated in the nucleus after 2 h, independent of whether an endocytosis or acidification inhibitor was present (Fig. 3G-H and S6B), placing the step requiring endocytosis between S2 and S3 cleavage. This sensitivity extends to other members of the Notch receptor family, as NICD1 generation was also reduced in SVG-A cells upon the inhibition of endocytosis or vesicular acidification after GI25X washout (Fig. S7A). Moreover, this sensitivity is cell line independent, as inhibitors of endocytosis or vesicular acidification reduced the generation of both NICD1 and NICD2 in 293T cells (Fig. S7B), U2OS cells (Fig. S7C), HeLa cells (Fig. S7D), and U251 cells (Fig. S7E). These results argue that bound ligand induces metalloprotease cleavage at S2 at the cell surface, and that the S2-cleaved form of Notch2 is then internalized to an intracellular compartment where it is cleaved by gamma secretase, thereby allowing NICD2 to access the nucleus.

### Passage of NICD2 from membrane to nucleus

In our dataset we identified a class of proteins whose relative abundance increases shortly after GSI washout (2 - 5 minutes) (Fig. 1C-D). Among the proteins in this cluster, we detected the ERM proteins Ezrin (EZR), Radixin (RDX), and Moesin (MSN), which play a role in linking membranes to the actin cytoskeleton (*45, 46*) (Fig. 4A-B). In addition to their architectural role, ERM proteins have been implicated in the maturation of endosomes and trafficking of EGFR (*47*). These results indicate that a pool of NICD2 molecules relocates to an ERM enriched microenvironment upon or immediately after gamma secretase cleavage to generate NICD2.

**Fig. 4.**
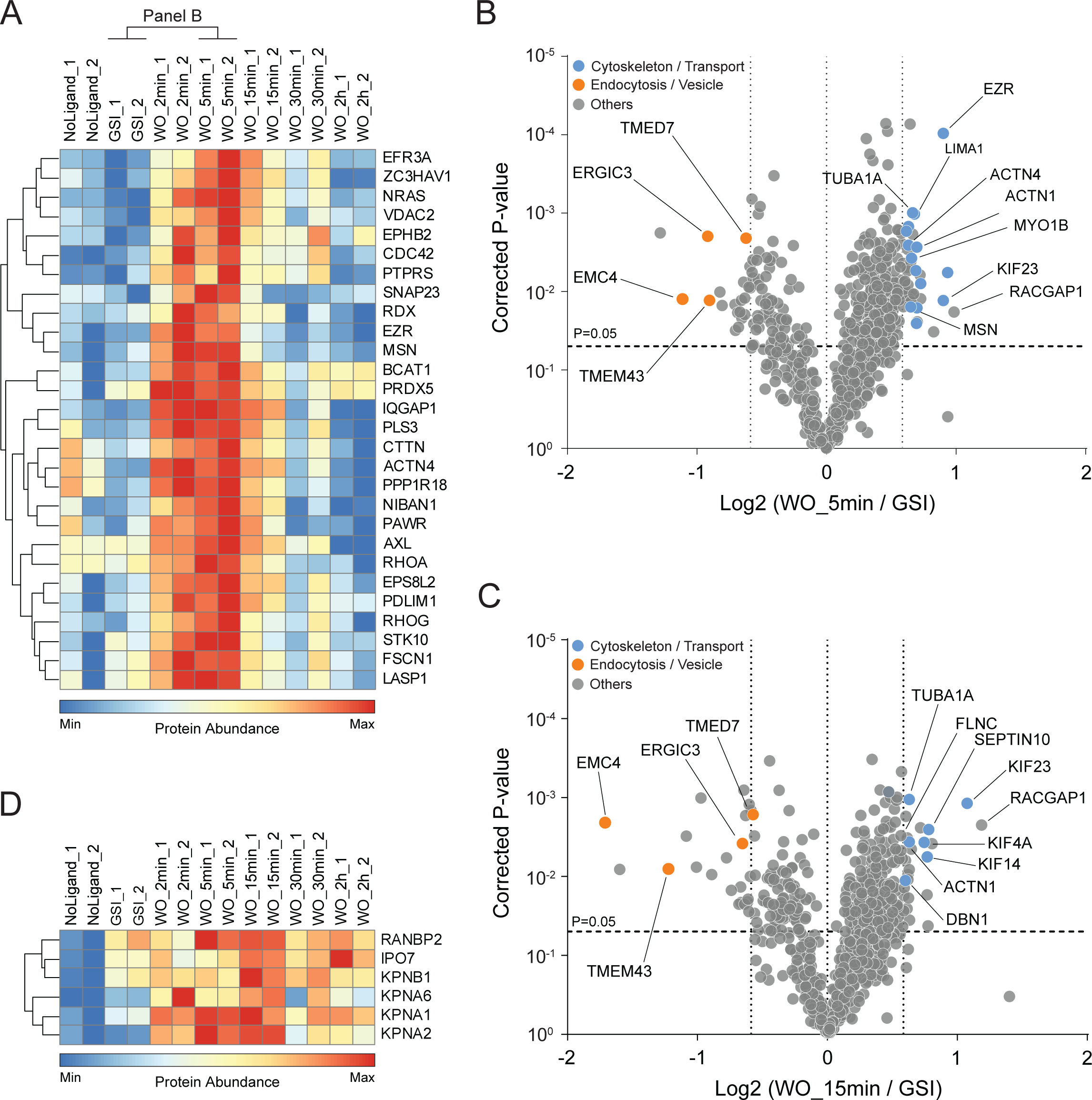
NICD2 microenvironment early after GSI washout. (**A**) Hierarchical clustering of the proteins characterized by a peak of relative abundance between 2 - 5 min after GSI removal, focusing on neighbors closest to Ezrin (EZR). (B,C) Volcano plots comparing Notch2 proximity-labeled proteins enriched at the 5 (**B**) and 15 (**C**) min timepoints after GSI removal when compared to GSI (t=0). Proteins related to actin, myosin, or cytoskeletal transport are labeled in blue, and proteins related to endocytosis or vesicular-mediated transport are labeled in orange. P-values are Benjamini-Hochberg corrected (p-value ≤ 0.05, FC ≥ 1.5). (**D**) Heatmap showing the enrichment pattern of proteins related to nuclear import identified by Notch2 proximity labeling.

To examine whether the activity of ERM proteins modulates Notch2 signaling in these cells, we treated cells either with control siRNA (siCtrl) or with siRNA that suppressed the expression of all three ERM proteins simultaneously (Fig. S8A-B), and activated signaling by GSI washout after culturing Notch2-Halo cells on immobilized Jag1. Both control-treated cells and cells treated with the ERM siRNA mixture exhibited a comparable increase in the amount of nuclear NICD2 2 h after washout (Fig. S8C-D). Likewise, control-treated cells and cells treated with the ERM siRNA mixture exhibited similar increases in the amount of NICD2 detectable by Western blot (Fig. S8E-H), and there was no significant difference between the two treatments when the induction of reporter gene activity was measured after inhibitor removal (Fig. S8I-K). In a complementary experiment, we also tested whether treatment with a dominant negative form of Ezrin (*48, 49*) affected nuclear accumulation of NICD2, and again did not observe a substantial difference among the vehicle-treated, wild-type Ezrin treated, and dominant-negative Ezrin-treated cells (Fig. S8L-M). Together, these results suggest that the ERM proteins are a marker of the compartment traversed by Notch2 shortly after GSI washout, but that they do not themselves substantially modulate Notch2 signaling in these cells.

At the 15 min time point after inhibitor washout, the enrichment of ERM proteins has abated and there is a period when relatively few proteins show an enrichment in labeling greater than two-fold when compared to baseline labeling in the presence of GSI (Fig. 4C). Although an increase in labeling of motor and cytoskeletal proteins is observed, this effect is small. The lack of strong enrichment of any particular motor protein or complex suggests that NICD2 is primarily cytoplasmic and is not actively transported through the cytosol (*i.e*. is not bound to and transported by a motor protein) to the nuclear membrane after liberation by gamma secretase. In the 5-15 min time window, the enrichment of proteins that participate in nuclear import of cargo (*50, 51*) (Fig. 4D), including the importin-beta subunit KPNB1 (*52*) and its associated adaptor Importin-7, as well as subunits of importin-alpha (KPNA1, KNPA2, and KPNA6) and the nuclear pore protein RANBP2 (also called Nup358) suggest that the nuclear import machinery facilitates the NICD2 nuclear entry step. This interpretation is consistent with previous work implicating nuclear import in facilitating the nuclear entry step for human Notch1 (*53*) and *Drosophila* Notch (*54*).

### NICD2 is nuclear and is associated with active transcription within 2 h

Several proteins exhibited a late labeling pattern with a strong peak enrichment at 2 h (Fig. 5A-B). MAML1, a component of Notch transcriptional complexes (*29, 30, 55*), was the most significantly enriched protein at this time point when compared to the baseline GSI condition (Fig. 5A), indicative of NICD2 nuclear entry (Fig. 5A). Analysis of the kinetics of MAML1 enrichment showed that increased labeling began at 30 min and was maximal by the 2 h timepoint (Fig. 1E).

**Fig. 5.**
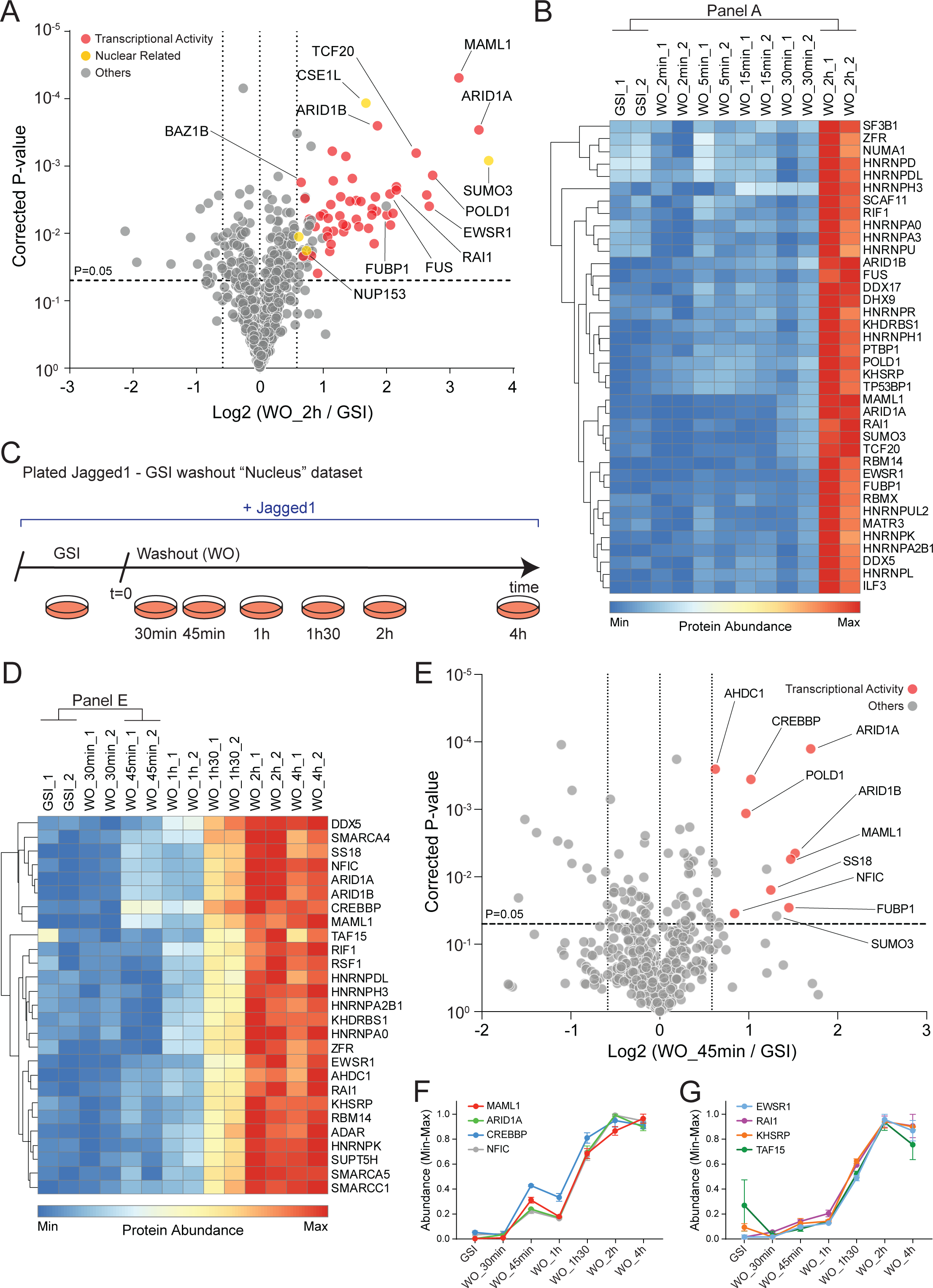
Nuclear accumulation of NICD2 and engagement with transcriptional regulators. (**A**) Volcano plot highlighting Notch2 proximity-labeled proteins enriched 2 h after GSI removal when compared to GSI (t=0). Significantly enriched proteins (p-value ≤ 0.05, FC ≥ 1.5) implicated in transcriptional activity are red, and other nuclear proteins are indicated in yellow. P-values are Benjamini-Hochberg corrected. (**B**) Heatmap showing kinetic profiles of proteins that cluster adjacent to MAML1 with strong enrichment 2 h after GSI washout. (**C**) Schematic showing the design for focused proximity labeling around the time of NICD2 nuclear entry using plated Jag1 as ligand and washout of GSI at time t=0 (see Fig. S6). (**D**) Heatmap of hierarchical clustering centered on proteins with kinetic profiles most closely related to MAML1 in the nuclear-centered proximity labeling dataset (see fig. S6). (**E**) Volcano plot of Notch2 proximity-labeled proteins enriched 45 min after GSI removal when compared to GSI (t=0) in the nuclear-centered dataset. Significantly enriched proteins (p-value ≤ 0.05, FC ≥ 1.5) implicated in transcriptional activity are red. P-values are Benjamini-Hochberg corrected. (**F**) Line plots showing the kinetic profiles of MAML1, the SWI/SNF chromatin-remodeling complex component ARID1A, CREBBP/p300, and the nuclear factor 1 C-type (NFIC) and (**G**) line plots showing the kinetic profiles of the transcriptional regulators EWSR1, RAI1, KHSRP, and TAF15 in the nuclear-centered dataset.

Strikingly, most of the other proteins showing significant, robust enrichment by 2 h are also implicated in transcription regulation or chromatin modification or remodeling (Fig. 5A-B). Hierarchical clustering revealed that the pattern of enrichment seen for MAML1 is shared by other transcriptional regulators, including RAI1, TCF20, and FUBP1 (Fig. 5B). Likewise, ARID1A and ARID1B of the SWI/SNF chromatin remodeling complex exhibited the same dynamics of labeling enrichment in this experiment. A similar time course of enrichment was also observed for the proteins FUS and EWSR1, two FET proteins that enter nuclear condensates and can influence transcription, and for several HNRNP proteins implicated in the regulation of splicing (*56, 57*) (Fig. 5B). Analysis of qRT-PCR data for *HES1, MYC*, and *TRIB1* as a function of time after inhibitor removal shows that transcriptional induction of direct Notch target genes also occurs as early as 30 min after GSI washout (Fig. S9A-C). Together, these data indicate that NICD2 has initiated the induction of a transcriptional response as early as 30 min after GSI washout and that the response is robust within 2 h, consistent with other reports showing that dynamic Notch binding sites in the genome become loaded with NICD and other co-factors within 2 h (*56*–*59*).

To further refine the temporal sequence of association of nuclear factors with NICD2, we acquired a second proximity labeling dataset focusing on timepoints between 30 min and 4 h after GSI washout (Fig. 5C and S9D-E). In this experiment, MAML1, CREBBP/p300, the nuclear factor 1 C-type protein NFIC, and proteins of the BAF chromatin remodeling complex (Fig. 5D-F) show enrichment by the 45 min timepoint, reaching maximum enrichment at the 2 h and 4 h timepoints (see Data File S4 for full quantification data of all proteins identified in this dataset). Enrichment of other transcriptional regulators (*e.g.,* RAI1), HNRNPs and FET proteins begins shortly afterward BAF labeling begins, at 1 h (Fig. 5G), indicative of the presence of NICD2 at loci of active transcription by this timepoint, consistent with the transcriptional induction observed by qRT-PCR at Notch-responsive genes (Fig. S9A-C).

## DISCUSSION

The overarching goal of this work was to achieve high spatiotemporal resolution for the molecular neighborhood of Notch as it transits from membrane to nucleus after proteolytic activation, because there is no real-time dynamic analysis available for this process. Our overall approach was to use Notch2-APEX2 time-resolved proximity labeling coupled with quantitative multiplexed proteomics to track the molecular microenvironment of endogenous Notch2 as a function of time after ligand stimulation and washout of a gamma secretase inhibitor. This unbiased approach allowed us to measure dynamic changes in the proteins in proximity to Notch before and after cleavage by gamma secretase, investigate the path and mode of transport of activated Notch2 from the membrane to the nucleus, and define the nuclear microenvironment of NICD2 during transcriptional induction.

Because soluble ligands do not activate Notch, it was not possible to use acute addition of the ligand to initiate the signal. It was necessary, therefore, to use GSI washout to achieve the temporal resolution required for dynamic analysis of the Notch molecular neighborhood. Cells were incubated overnight on immobilized ligand in the presence of the GSI because the process of removing cells from a TC dish and transferring them to another dish triggers Notch proteolysis independent of ligand. By allowing the cells to incubate overnight on the immobilized ligand in the presence of GSI, we made sure that the Notch2 protein tracked in the proximity-labeled analysis would be in the NEXT form, with GSI washout synchronizing access to gamma secretase and enabling direct tracking of downstream steps as a function of time. While we certainly appreciate that accumulating Notch in the NEXT form is a minor limitation of the study, a particular strength of this design is that it enabled time-resolved analysis of proximity with Notch2 at natural abundance. The synchronization achieved with GSI removal allowed us to faithfully monitor the molecular neighborhood of Notch2 as a function of time after inhibitor removal, as judged by the observation that the enrichment of MAML1 in the molecular neighborhood of Notch2 only occurs 30 min or more after GSI removal. This timing of nuclear access of Notch2 is concordant with that seen in other studies (*11, 60*).

Several mechanistic findings emerged from these studies. First, the enrichment of proteins associated with vesicular transport and endocytosis after ligand exposure in GSI-treated cells suggested that gamma secretase cleavage of Notch2 occurs in an intracellular compartment. This conclusion was supported by additional experiments tracking gamma secretase cleavage activity after washout of inhibitors of ligand-dependent metalloprotease (S2) cleavage in the presence of inhibitors of endocytosis or vesicle acidification, which showed that S2-processed Notch2 must enter an intracellular compartment to be cleaved by gamma secretase. Whereas older studies have reached differing conclusions about whether gamma secretase processes substrates at the plasma membrane (*12, 13*) or in an intracellular compartment (*14*–*16*), our finding that gamma secretase cleavage of Notch2 occurs in an intramembrane compartment agrees with recent proteomics studies investigating APP cleavage using affinity capture of the early endosome-associated protein EEA, in which APP/Aβ cleavage products of gamma secretase accumulate in early/sorting endosomes (*61*).

Second, our data suggest that Notch2 molecules transiently pass through a microenvironment enriched in ERM proteins between 2-5 min after GSI washout. These proteins appear to identify a compartment traversed by Notch2 upon or immediately after gamma secretase cleavage to generate NICD2. In comparison, few proteins are enriched at the 15 and 30 min time points, consistent with the idea that NICD2 moves by passive diffusion to the nucleus after gamma secretase cleavage, with a possible preference for migration in proximity to or along actin filaments, before engaging the nuclear import machinery.

Third, we find that NICD2 enters a nuclear microenvironment enriched in components associated with a transcriptional response as early as 30-45 min after GSI washout and persists through the final 4 h timepoint. This timing of transcriptional induction is consistent with previous real-time luciferase complementation studies using ectopic Notch1 and RBPJ expression (*11*) and with the onset and duration of Notch-induced transcription in fly models and in cancer cells (*62*– *65*). In addition to MAML1, the nuclear proteins most rapidly recruited to NICD2 are CREBBP/p300, a well-established partner of MAML1 in NICD-dependent transcriptional induction (*58, 59, 66, 67*), and components of the BAF chromatin remodeling complex. Strikingly, the SWI/SNF chromatin remodeling complex is crucial to render enhancers responsive to Notch in *Drosophila* (*57*); the basis for recruitment of the BAF complex to Notch-responsive elements should be fertile ground for future study.

It is also notable that proteins implicated in NICD degradation were not identified in this study. Because the association of NICD2 with proteins that participate in its degradation may take place transiently and in catalytic amounts, it is possible that our studies were not sensitive enough to identify these interactions.

Overall, our proximity labeling studies and follow-up cellular assays serve as the basis for a well-defined spatiotemporal model of the pathway traversed by Notch upon proteolytic activation (Fig. 6). Ligand engagement first induces S2 cleavage of Notch at the cell surface, followed by entry of truncated Notch2 into an endocytic compartment for cleavage by gamma secretase. As early as 30 min after gamma secretase cleavage, NICD enters the nucleus and by 45 min has begun to recruit CREBBP/p300 and chromatin remodeling complexes to initiate transcription of responsive genes, with evidence for recruitment of proteins involved in transcription-coupled splicing events after 60 – 90 min. More broadly, our work with Notch as a signaling protein of interest represents a proof-of-concept for future quantitative analyses of other signal transduction systems, showing that time-resolved proximity labeling with APEX2 combined with multiplexed proteomics can elucidate the temporal and spatial dynamics of endogenous proteins and the evolution of their microenvironments during signaling.

**Fig. 6.**
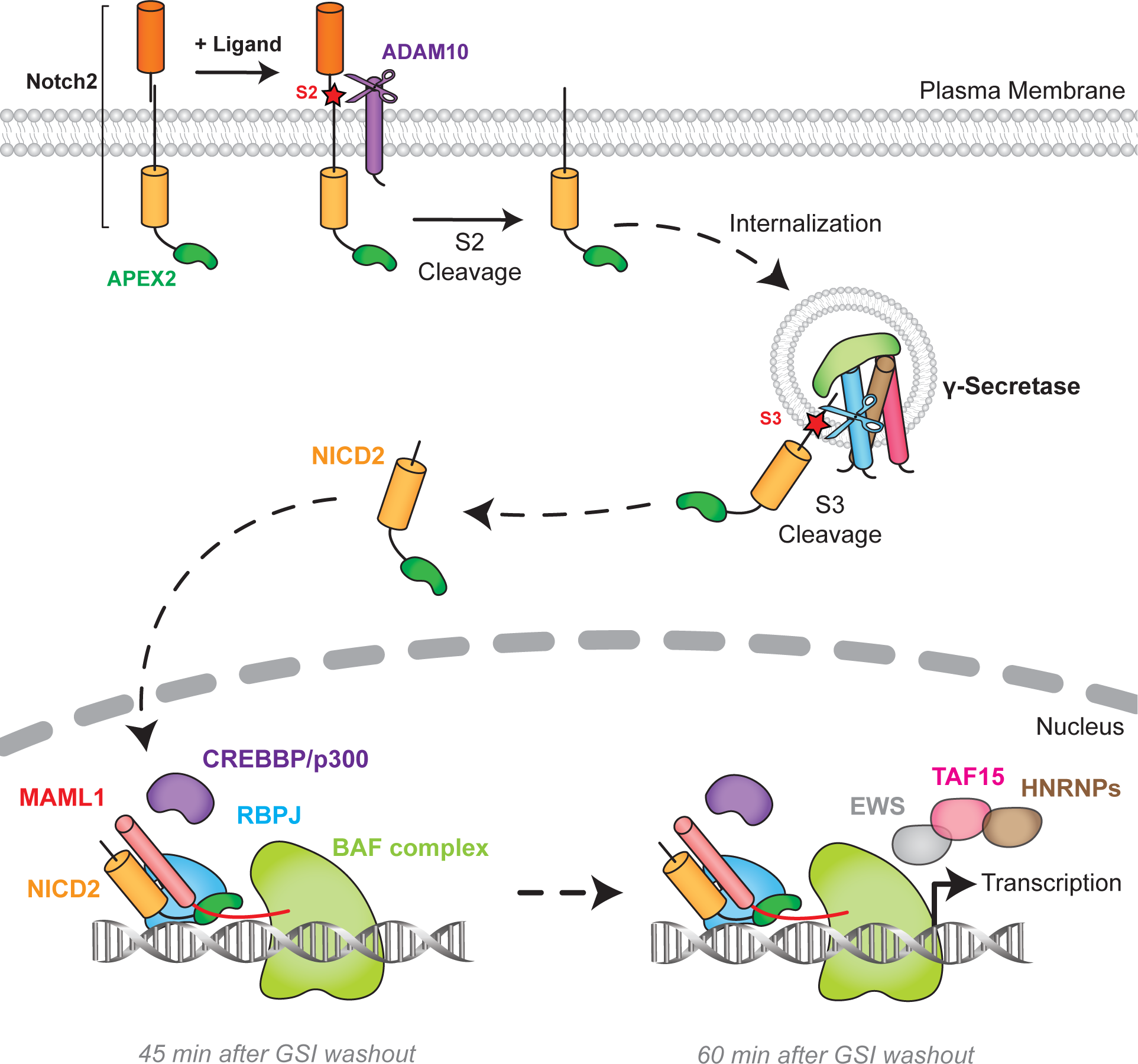
Model including Notch2 internalization as a mechanistic step in Notch activation and signaling. Upon ligand stimulation of SVG-A cells, Notch2 is cleaved at site S2 by ADAM10, followed by entry of S2 processed Notch2 molecules into an intracellular compartment. Internalized Notch2 molecules are then cleaved at site S3 by gamma secretase, generating NICD2, which access the nucleus about 30-45 min after GSI washout and induces its transcriptional response.

## MATERIALS & METHODS

### Cell line generation, cultivation, and manipulation

All cell lines were maintained in DMEM with L-glutamine (Corning) supplemented with 10% fetal bovine serum (FBS, Gemini Bio-Sciences) and 1% penicillin-streptomycin (Gibco) at 37°C and 5% CO_2_. Cell lines were tested for mycoplasma on a regular basis. CRISPR/Cas9 gene editing was used to knock out Notch2 in SVG-A cells. For the SVG-A Notch2 knockout cell line, a pX459 plasmid containing a gene-specific guide RNA (gRNA) was transfected using Lipofectamine 2000 (ThermoFisher Scientific) according to the manufacturer’s instructions (see Table S2 for gRNA sequences used in this study). 48 h after transfection, cells were selected using 2 µg/mL puromycin for 3 days, and single cells were then isolated by flow cytometry using a BD FACSAria cell sorter. Knockout clones were identified by DNA sequencing after PCR amplification of genomic DNA at the mutated locus, and the loss of protein expression was confirmed by Western blotting.

CRISPR/Cas9 genome editing was also used to fuse an APEX2-HA tag at the C-terminus of endogenous Notch2 in SVG-A cells. SVG-A cells were co-transfected with a pX459 plasmid containing gRNA targeting Notch2 (see Table S2 for gRNA sequences) and a pUC19 donor plasmid containing a GGAG linker-APEX2-HA-T2A-mNeonGreen cassette flanked by Notch2 genomic locus homology arms each approximately one kilobase in length. Seven days after transfection, single cells expressing mNeonGreen were isolated by FACS using a Sony SH800S cell sorter, and individual clones were expanded in 96 well plates. Confirmation of successful tagging and identification of homozygous clones was carried out by PCR amplifying the region of the insertion with flanking primers outside of the genomic region covered by the homology arms, followed by Sanger DNA sequencing for the positive homozygous clones. Clones were further evaluated to assess the amount of expressed Notch2 protein by Western blotting, the amount of surface staining by flow cytometry on a BD Accuri C6 Plus flow cytometer, Notch2 transcriptional activity using a luciferase reporter assay (described below), and APEX2-dependent protein biotinylation by Western blotting using Streptactin-HRP (Bio-Rad). A similar strategy and gRNA were used to insert a HaloTag at the C-terminus of endogenous Notch2.

### RNA extraction, reverse transcription, and qRT-PCR

RNA was harvested from confluent wells of a 12 well plate and isolated using TRIzol (Ambion) according to the manufacturer’s instructions. All samples were DNase-treated by incubating the extracts with DNase I recombinant (Roche) at 37°C for 30 minutes. After RNA was recovered from sequential 24:1 chloroform:isoamyl alcohol (Sigma-Aldrich) and ethanol precipitations, reverse transcription was performed using the iScript kit (Bio-Rad) according to the manufacturer’s instructions with 250 ng of purified RNA as template. qPCR was done in a 10 µL reaction mix with 0.25 µM forward and reverse primers using the PowerUp SYBR green master mix (ThermoFisher Scientific) with standard thermocycler parameters on a BioRad CFX384 qPCR instrument. A melting curve step right after PCR amplification was performed to confirm primer specificity. Three technical replicates for each of 3 biological replicates were analyzed. The primer sequences are listed in Table S3. Gene expression was normalized to GAPDH for each condition. For the time-resolved GSI washout experiments presented in Fig. S9A-C, the data were normalized to the control condition (No Ligand conditions in GSI washout), which was assigned a value of 1.

### Western blotting

Adherent cells were washed in ice-cold Dulbecco’s Phosphate Buffered Saline (DPBS) and lysed in gel-loading buffer (2% SDS, 60 mM Tris–HCl pH 6.8, 100 mM DTT, 10% glycerol, and 0.005% bromophenol blue), scraped off the plate, boiled at 95°C for 5 minutes, and subjected to SDS-PAGE. Proteins were then transferred to a PROTRAN 0.2 µm nitrocellulose membrane (Cytiva) and stained with Ponceau S (Sigma-Aldrich). Membranes were incubated in 5% (w/v) non-fat dry milk in TBST (20 mM Tris, 150 mM NaCl, 0.2% Tween-20, pH 7.6) at room temperature for at least 1 h. Blocked membranes were incubated with various primary antibodies, or with a Streptactin-HRP (BioRad), diluted in TBST supplemented with 5% non-fat dry milk overnight at 4°C with gentle shaking. Membranes were washed 3 times with TBST at room temperature and incubated with appropriate secondary antibodies for 1 h at room temperature with gentle shaking. Blots were washed 3 times with TBST and imaged using an Odyssey Infrared Imaging System (LI-COR Biosciences) for IRDye-conjugated secondary antibodies or on a Chemidoc (Bio-Rad) using an Amersham ECL Western Blotting Detection kit (GE Healthcare) for Streptactin-HRP. See Table S1 for a list and working conditions of the antibodies used in this study.

### Recombinant Jag1-Fc expression and purification

The extracellular domain (ECD) of human Jag1 (aa 1-1067) was fused to the Fc region (CH2 and CH3 domains) and hinge region of the human IgG1 heavy chain in the pFUSE-Fc1 vector (InvivoGen). Jag1ECD-Fc protein was expressed in Expi293F cells grown in Expi293F expression medium at 37°C in an 8% CO_2_ incubator with constant shaking. Cells were grown to a density of 3×10^6^ cells/mL in a final volume of 1 L and transiently transfected using FectoPro transfection reagent (Poly-plus) with 1 mg of purified plasmid at a 2:1 DNA/FectoPro ratio. 22 h after transfection, 5 mM Valproic acid sodium salt (Sigma-Aldrich) and 10 mL of 45% D-(+)-Glucose solution (Sigma-Aldrich) were added. After 7 days of culture, the media supernatant was collected after removal of debris by centrifugation at 4,000 xg for 15 min at 4°C followed by a filtration step. Filtered media was then loaded onto a Protein A (Millipore) column prewashed in ice cold HEPES-buffered saline (HBS) buffer (20 mM HEPES pH 7.3, 150 mM NaCl). Bound protein was eluted in 100 mM glycine, pH 3.0 and neutralized with 1 M Tris buffer pH 7.3. Eluted protein was buffer exchanged and concentrated in HBS. Protein purity was assessed by separation on SDS-PAGE after staining with SafeBlue (ThermoFisher Scientific). The purified protein was diluted to a final concentration of 200 µg/mL in HBS supplemented with 10% glycerol, aliquoted, flash frozen, and stored at −80°C.

### Activation of Notch2 by immobilized Jag1-Fc

Recombinant Jag1-Fc was immobilized by overnight incubation at 4°C in individual wells of non-tissue culture-treated 6 or 12 well plates (VWR) at a final concentration of 2 µg/mL in DPBS containing 10 µg/mL poly-D-lysine (ThermoFisher Scientific). For imaging studies, the ligand was immobilized in 24 well plates containing pre-washed glass coverslips overnight. The next day, the Jag1-Fc and poly-D-lysine mixture was removed and the cells were added to the coverslips and incubated as indicated.

### Luciferase reporter assays

SVG-A cells were transfected with a mixture of TP1-firefly luciferase and pRL-TK (Promega) plasmids at a 49:1 ratio using Lipofectamine 2000 (ThermoFisher Scientific) according to the manufacturer’s instructions. Culture media was replaced 4 h after transfection and the cells were incubated overnight. The next day, cells were detached with 0.5 mM EDTA, recovered by centrifugation, and added to plates pre-coated with recombinant Jag1-Fc in media containing 100 nM Compound E (GSI) or 5 µM GI254023X. At that time, luciferase assays were performed using a Dual-Luciferase reporter assay system (Promega) according to the manufacturer’s instructions. For experiments investigating the effects of endocytosis or vesicular acidification, cells were preincubated with hydroxy-dynasore, bafilomycinA1, or chloroquine for 1 h before removal of GI254023X, and cells were harvested 6 hours later. Luminescence was measured on a GloMax plate reader (Promega). Three technical measurements were performed for each of three biological replicates. The ratio of firefly to Renilla luminescence was calculated and normalized to the control condition (presence of GI254023X and DMSO) and assigned a value of 100 percent.

### Flow Cytometry

Cells were washed with DPBS, detached using 0.5 mM EDTA in DPBS for 5 minutes at room temperature, pelleted, and washed again with DPBS. For permeabilized conditions, cells were fixed in 0.01% Formaldehyde in DPBS for 15 minutes at room temperature, washed, then incubated in 0.5% (v/v) Tween-20 in DPBS for 5 minutes at room temperature, and finally washed once again. Cells were then incubated for 45 minutes at 4°C in the dark with 2 µL of the appropriate antibody diluted in 2% (v/v) FBS in DPBS. Cells were then washed twice with DPBS and analyzed on a BD Accuri C6 Plus flow cytometer.

### APEX2 proximity labeling

SVG-A Notch2-APEX2-HA cells were incubated overnight on tissue culture dishes in the presence of immobilized Jag1 or with Jag1-expressing A673 cells in the presence of 100 nM GSI or 10 µM BB94. The next day, cells were washed three times with DMEM to remove the inhibitor. The time of addition of fresh media after the last wash was set to t=0 in defining the timepoints analyzed in each experiment. For each timepoint, the media was exchanged with fresh media containing an additional 2 mM biotin phenol (BP, Iris Biotech Gmbh, LS-3500) exactly 1 h before 0.1 mM hydrogen peroxide (H_2_O_2_, Sigma-Aldrich) was added to initiate the labeling reaction. Immediately after adding H_2_O_2_, the culture dishes were gently rocked several times to ensure optimal H_2_O_2_ distribution. Exactly 1 min after the addition of H_2_O_2_, the media was quickly aspirated and cells were washed three times with quenching buffer (DPBS supplemented with 10 mM sodium ascorbate, 5 mM Trolox [Sigma-Aldrich], and 10 mM sodium azide). Cells were then scraped in quenching buffer, harvested by centrifugation, and cell pellets were flash-frozen and stored at −80°C until streptavidin pull-down was performed. An H_2_O_2_ stock solution (30%, v/v) was freshly diluted to 1 M in DPBS immediately before each experiment. See Table S1 for a list of key reagents used for the proximity labeling.

### Streptavidin pull-down

All solutions and buffers were freshly prepared and filtered. Streptavidin capture of biotinylated proteins was performed as previously described (*22, 68*). Briefly, frozen cell pellets were lysed in ice cold lysis buffer (8 M Urea, 100 mM sodium phosphate pH 8.0, 1% SDS (w/v), 100 mM NH_4_HCO_3_, 10 mM TCEP) and pipetted repeatedly on ice to ensure proper cell lysis. Lysates were then homogenized by passing them through QIAshredder cartridges (Qiagen). Proteins were precipitated by adding an equal volume of ice cold 55% Trichloroacetic acid (TCA, Sigma-Aldrich), incubated 15 min on ice, and then pelleted by centrifugation at 21,000 x g at 4°C for 10 min. The protein pellet was washed with −20°C cold acetone (Sigma-Aldrich), vortexed, and centrifuged at 21,000 x g at 4°C for 10 min. Following centrifugation, acetone was removed and the pellet was washed with acetone 3 more times. After the last wash, the pellet was resuspended in lysis buffer as described above, vortexed, and rotated at room temperature until fully dissolved, allowing reduction of proteins by TCEP at the same time.

Resuspended proteins were centrifuged at 21,000 x g at room temperature for 10 min and the clear supernatant was transferred to a new microcentrifuge tube. To alkylate free cysteines, freshly prepared 400 mM iodoacetamide stock solution (Sigma-Aldrich) in 50 mM ammonium bicarbonate was added to the supernatant at a final concentration of 20 mM, and the samples were immediately vortexed and then incubated in the dark at room temperature for 25 min. After alkylation, freshly prepared dithiothreitol (DTT, Sigma-Aldrich) was added to a final concentration of 50 mM to quench the reaction. Finally, water was added to each sample to reach a final concentration of 4 M urea and 0.5% (w/v) of SDS.

125 µL of streptavidin magnetic bead suspension (ThermoFisher Scientific) was washed twice with 4 M urea, 0.5% SDS (w/v), 100 mM sodium phosphate pH 8.0 and added to each sample. The tubes were gently rotated overnight at 4°C. Following capture of biotinylated proteins, the magnetic beads were washed 3 times with 4 M urea, 0.5% SDS (w/v), 100 mM sodium phosphate pH 8.0, 3 more times with the same buffer without SDS, and finally 3 more times with DPBS. The beads were transferred to new tubes for each change of wash buffer. See Table S1 for a list of key reagents used for the streptavidin enrichment of biotinylated proteins.

### On-beads digestion and tandem mass tag (TMT) labeling

The streptavidin beads were subjected to on-bead protease digestion in 50 µl digestion buffer (200 mM EPPS pH 8.5 with 2% acetonitrile [v/v]) along with LysC (Wako) at an enzyme-to-substrate ratio of 1:50. The samples were incubated at 37°C for 3 h. Then 50 µl of digestion buffer with trypsin (Promega) was added at an enzyme-to-substrate ratio of 1:100. The digestion was continued at 37°C overnight with gentle agitation. The clear supernatants of digested protein were separated from beads with a magnetic rack and transferred to fresh tubes.

For the TMT reaction, 30% acetonitrile (v/v) was added to the digested protein and then labeled using a TMT isobaric mass tagging kit (ThermoFisher Scientific). The TMT reaction was performed for 1 h according to the manufacturer’s instructions. TMT labeling efficiency and ratios were measured by LC-MS3 analysis after combining equal volumes from each sample. Once the labeling efficiency was determined to be >95%, the TMT reactions were quenched with hydroxylamine 0.5% v/v for 15 min and acidified with formic acid. Samples were then pooled and dried to near completion under reduced pressure before resuspension in 1% formic acid and fractionation using a Pierce High pH Reversed Phase Peptide Fractionation Kit (ThermoFisher Scientific) with modified elution of 12 sequential fractions (10%, 12.5%, 15%, 17.5%, 20%, 25%, 30%, 35%, 40%, 50%, 65% and 80% acetonitrile). Fractions were then combined into pairs as follows, 1+7, 2+8, 3+9, 4+10, 5+11, 6+12, to give the final six fractionated samples. The resulting fractions were dried under reduced pressure and then desalted using a stage tip protocol (*69*).

### Mass spectrometry acquisition and data analysis

Data were acquired on an Orbitrap Fusion Lumos instrument (ThermoFisher Scientific) coupled to a Proxeon Easy-nLC 1200 UHPLC. Peptides were injected onto a 100 µm (inner diameter) capillary column (∼30 cm) packed in-house with C18 resin (2.6 µm, 150Å, ThermoFisher Scientific). Peptide fractions were separated with a 4 h acidic acetonitrile gradient from 5-35% Buffer B (Buffer A = 0.125% formic acid, Buffer B = 95% acetonitrile, 0.125% formic acid). All data were collected with a multi notch MS3 method (*70*). MS1 scans (Orbitrap analysis; resolution 120,000; mass range 400–1400 Th) were followed by MS2 analysis with collision-induced dissociation (CID, CE=35) and a maximum ion injection time of up to 120 ms and an isolation window of 0.4 m/z, using rapid scan mode. To obtain quantitative information, MS3 precursors were fragmented by high-energy collision-induced dissociation (HCD, CE=65) and analyzed in the Orbitrap at a resolution of 50,000 at 200 Th with max injection time set to 650 ms. Raw spectra were converted to mzXML to correct monoisotopic m/z measurements and to perform a post-search calibration.

Spectra were searched using SEQUEST (v.28, rev.12) software against the UniProt human reference proteome (downloaded 02-25-2020), containing common contaminants and reversed order protein sequences as decoy hits (*71*). Searches were performed with a precursor mass tolerance of 20 ppm, and the fragment-ion tolerance was set to 0.9 Da. For searches a maximum of 2 missed trypsin cleavage sites were allowed. Methionine oxidation (+15.9949 Da) was set as a variable modification, while cysteine carboxyamidomethylation (+57.0215) and TMT (+229.1629) or TMT16 (+304.2071 Da) tags on lysine and peptide N-termini were set as a static modification. Peptide spectral matches (PSM) were filtered by linear discriminant analysis (LDA), using a target-decoy database search to adjust the PSM false discovery rate to 1% and protein level FDR of 1% (*72*). For MS3 relative quantification, peptides were filtered for an MS2 isolation specificity of >70%, and a total TMT summed signal to noise of >200 for all channels in the multiplex. Further details of the TMT quantification method and search parameters applied were described previously (*73*).

Proteomics raw data and search results were deposited in the PRIDE archive for each multiplex experiment with accession number: PXD039008 (immobilized ligand with GSI experiment, Data File S1), PXD042123 (co-culture with Jag1-expressing A673 cells, Data Files S2 & S3), and PXD039010 (immobilized ligand with GSI nuclear centered experiment, Data File S4).

### Dynasore-OH, bafilomycin A1, and chloroquine treatments

For immunofluorescence experiments, Western blots, and reporter gene assays using dynasore-OH (Sigma-Aldrich), bafilomycinA1 (Selleck Chemicals), or chloroquine (Sigma-Aldrich), cells were incubated overnight on immobilized Jag1 in the presence of GSI or GI254023X. The next day, dynasore-OH (Sigma-Aldrich), bafilomycinA1, or chloroquine was added to the media and pre-incubated with the culture for 1 h before removing the GSI or GI254023X. Cells were maintained in the continued presence of dynasore-OH, bafilomycinA1, or chloroquine, at 37°C for the indicated time before analysis. Dynasore-OH incubation was performed in serum-free DMEM.

### Immunofluorescence and image processing

SVG-A Notch2-HaloTag cells were grown on ligand-coated coverslips. Cells were labeled with JaneliaFluorX549 HaloTag ligand (a gift from Luke Lavis, Janelia Research Campus) at a final concentration of 100 nM in media for 15 min at 37°C. The media was then removed, cells washed with fresh media, and returned to the incubator for 1 h to allow newly synthesized labeled Notch2 to be delivered to the plasma membrane. Cells were treated with the indicated vesicular/transport inhibitors prior to GI254023X/GSI washout, as described above.

2 h after washout of GI254023X or GSI, cells were washed 3 times in DPBS, fixed with 4% paraformaldehyde (PFA, Sigma-Aldrich) for 15 min at room temperature (RT), washed three times in DPBS, and quenched with 0.1 M Glycine pH 7.5 in DPBS for 15 min at RT. After another three PBS washes, fixed cells were permeabilized with 0.1% Triton X-100 in DPBS for 10 min at room temperature followed by three washes in DPBS and blocking in 5% BSA (w/v) in DPBS for 1 hour at room temperature. Cells were then incubated for 1 h with primary antibodies diluted in blocking buffer at room temperature. After three washes in DPBS, the cells were incubated with secondary antibody (Alexa Fluor Plus 647-conjugated anti-rabbit, ThermoFisher Scientific A32795) diluted in blocking buffer for 45 min at room temperature followed by three washes in DPBS. For DNA staining, cells were incubated with SYTOX Green Nucleic Acid Stain (ThermoFisher Scientific) according to manufacturer’s recommendations followed by three washes in DPBS. Coverslips were then mounted with ProLong Gold Antifade Mountant with DAPI (ThermoFisher Scientific) or without DAPI if already labeled with SYTOX Green. Coverslips were stored at 4°C before image acquisition.

Images were acquired using a Marianas system (Intelligent Imaging Innovation) composed of a Zeiss Axio-Observer Z1 (Carl Zeiss) equipped with a 63x objective (Plan-Apochromat, NA 1.4, Carl Zeiss), a spinning disk confocal head (CSU-XI, Yokogawa Electric Corporation) and a spherical aberration correction system (Infinity Photo-Optical). Excitation light was provided by 405, 488, 561, or 640 nm solid-state lasers (Sapphire, 50 mW, Coherent Inc) coupled to an acoustic-optical tunable filter. Laser power and exposure times were kept the same for all experiments. Z stacks of 38 x-y confocal images were obtained in 270 nm z-steps using a cooled QuantEM^TM^ 512SC CCD camera (Photometrics). Images presented in Fig. S6 (effect of ERM proteins on Notch2 signaling) were acquired using a Prime 95B Scientific CMOS camera (Photometrics), giving different signal arbitrary units from the ones recorded with the QuantEM^TM^ 512SC CCD camera used in the other experiments. All equipment was controlled by SlideBook acquisition software (Intelligent Imaging Innovations). Image processing was performed using Fiji software (*74*). For nuclear intensity measurements, the nuclei were identified by DAPI staining and the Mean Fluorescence Intensity (MFI) of each indicated channel was measured by applying a nucleus mask.(*74*)

### siRNA-mediated knockdown of ERM proteins and evaluation of dominant-negative Ezrin

Silencer^©^ Select siRNAs for Ezrin (s14795), Radixin (s11899), Moesin (s8984), and the siCtrl AM4611 were all purchased from ThermoFisher Scientific. For siRNA-mediated knockdown of ERM (Ezrin, Radixin, Moesin) proteins, the cells were reverse transfected with the indicated siRNAs using Lipofectamine RNAiMAX transfection reagent according to manufacturer’s recommendations. Parental or Notch2-HaloTag SVG-A cells were seeded in a 6-well plate along with the transfection mix containing 3 µL of Lipofectamine RNAiMAX, and a total of 30 pmole of siRNA in 500 µL of OptiMEM. A portion of each sample (siCtrl or siERM) from each experiment (immunofluorescence, NICD2 Western blotting, and luciferase reporter assay) was set aside, lysed in gel-loading buffer as already described, and subjected to western blotting against Ezrin, Radixin, and Moesin to evaluate depletion efficiency. Immunofluorescence analyses and Western blots were performed at 0 (GSI) or 2 h (WO_2h) after GSI washout as described above. For luciferase reporter experiments, the cells were transfected the next day with a mixture of TP1-firefly luciferase and pRL-TK (Promega) plasmids at a 49:1 ratio using Lipofectamine 2000 (ThermoFisher Scientific) and read out as described above 24 h later.

For Ezrin dominant negative experiments, the full-length human Ezrin sequence, or a sequence comprising the amino acids 1-300, was inserted N-terminal to mNeongreen in pcDNA3.1. SVG-A Notch2-HaloTag cells were transfected in 6 well plate format with 2 µg of DNA per well using Lipofectamine 2000 (ThermoFisher Scientific) according to the manufacturer’s instructions. Culture media was replaced 4 h after transfection. The next day, the transfected cells were detached and plated on ligand-coated coverslips in the presence of 100 nM GSI and incubated overnight. The following day, the GSI was removed, and the cells fixed and processed for immunofluorescence as described above.

### Statistical analysis

Statistical analyses of fluorescence microscopy, luciferase reporter assays, qRT-PCR, western blotting, and flow cytometry experiments were performed using GraphPad Prism software (GraphPad). Two-way ANOVA followed by Tukey’s pairwise comparison was used for statistical comparisons. All error bars denote mean ± standard deviation (mean ± SD). *p ≤ 0.05, **p ≤ 0.01, ***p ≤ 0.001. The number of individual experiments analyzed is indicated in the figure legends.

## SUPPLEMENTARY MATERIALS

Figs. S1 to S9

Table S1: List of key reagents used in this study.

Table S2: List of CRISPR reagents used in this study.

Table S3: List and sequence of qRT-PCR primers used in this study.

Data File S1. List of proteins identified and their enrichment levels for Notch2-APEX2 using immobilized ligand with GSI washout.

Data File S2. List of proteins identified and their enrichment levels for Notch2-APEX2 using co-culture with A673 cells - Broad.

Data table S3. List of proteins identified and their enrichment levels for Notch2-APEX2 using co-culture with A673 cells - Early.

Data File S4. List of proteins identified and their enrichment levels for Notch2-APEX2 using immobilized ligand with GSI nuclear centered washout.

## ACKNOWLEDGMENTS

We thank members of the Blacklow lab for helpful discussions.

## AUTHOR CONTRIBUTIONS

A.P.M. and S.C.B. conceived the project. S.C.B. and T.K. acquired funding. A.P.M. performed and analyzed experiments. E.D.E. performed initial characterization of the parental SVG-A cell line and designed and generated Notch2-HaloTag knockin and Notch2 knockout lines. G.A.B., R.J.E., and M.K., processed and analyzed mass spectrometry data. L.T., G.S., and T.K., assisted with and gave technical advice for microscopy experiments. J.M.R., J.C.A., A.N.D., and M.K. assisted with data analysis and interpretation. A.P.M. and S.C.B. wrote the manuscript with input from all authors. All authors provided feedback and agreed on the final manuscript.

## DATA AND MATERIAL AVAILABILITY

Proteomics raw data and search results were deposited in the PRIDE archive for each multiplex experiment with accession number: PXD039008 (immobilized ligand with GSI experiment), PXD042123 (both the broad and early co-culture with Jag1-expressing A673 cells experiments), and PXD039010 (immobilized ligand with GSI nuclear centered experiment). All other data needed to evaluate the conclusions in the paper are present in the paper or the Supplementary Materials.

## Supplementary Materials

**Fig. S1.**
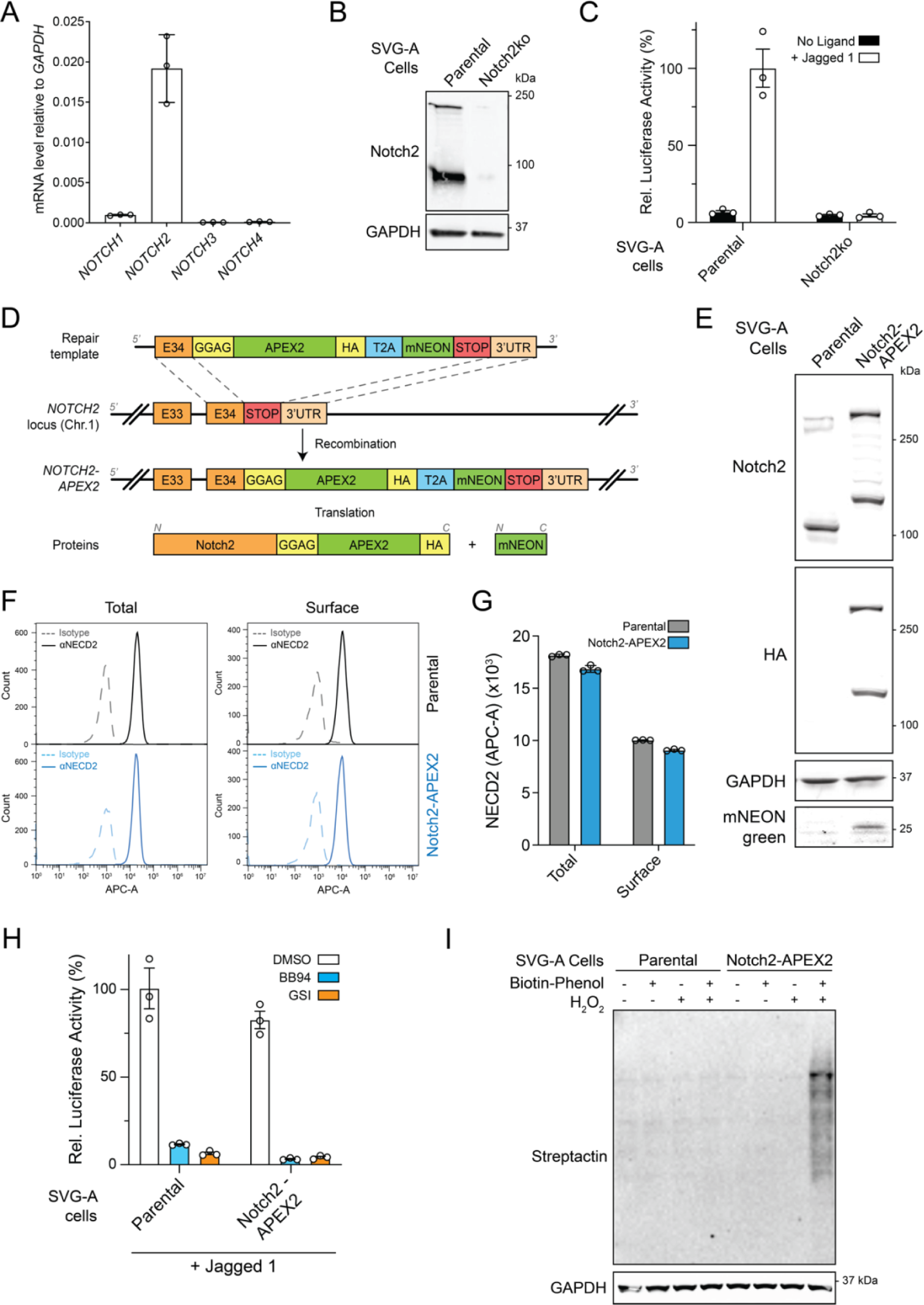
SVG-A Notch2-APEX2 cell line validation, related to Fig. 1. (A) qRT-PCR showing the relative mRNA amounts of the four human Notch proteins in SVG-A cells. (B) Western blot probing for Notch2 in lysates from parental and Notch2 knockout SVG-A cells. (C) Notch luciferase reporter assay, measuring relative luciferase activity in parental or Notch2 knockout SVG-A cells cultured with or without immobilized Jag1. (D) Design strategy for CRISPR/Cas9-mediated Notch2-APEX2 knock-in in SVG-A cells. An APEX2-HA-T2A-mNEONgreen cassette was inserted at the C-terminus of the genomic locus of the *NOTCH2* gene. The addition of the T2A-mNeonGreen enabled fluorescence-activated cell sorting (FACS) of clones with genomic integration of the repair template. (E) Western blot probing for Notch2 in lysates from parental and Notch2-APEX2 knock-in cells. (F) Representative flow cytometry analysis for total (permeabilized) or surface Notch2 in parental and Notch2-APEX2 knock-in SVG-A cells. (G) Quantification of the mean fluorescence intensity of surface Notch2 presented in panel F. (H) Notch luciferase reporter assay, measuring relative luciferase activity in parental or Notch2 knockout SVG-A cells cultured with or without immobilized JAG1 in the presence of DMSO carrier, 100 nM Compound E (GSI), or 10 µM BB94 (ADAM10 inhibitor). (I) Streptactin blot of parental and Notch2-APEX2 SVG-A cells probing biotinylation as a function of added biotin-phenol, H_2_O_2_, or both molecules. All quantifications presented in this figure are from three independent experiments.

**Fig. S2.**
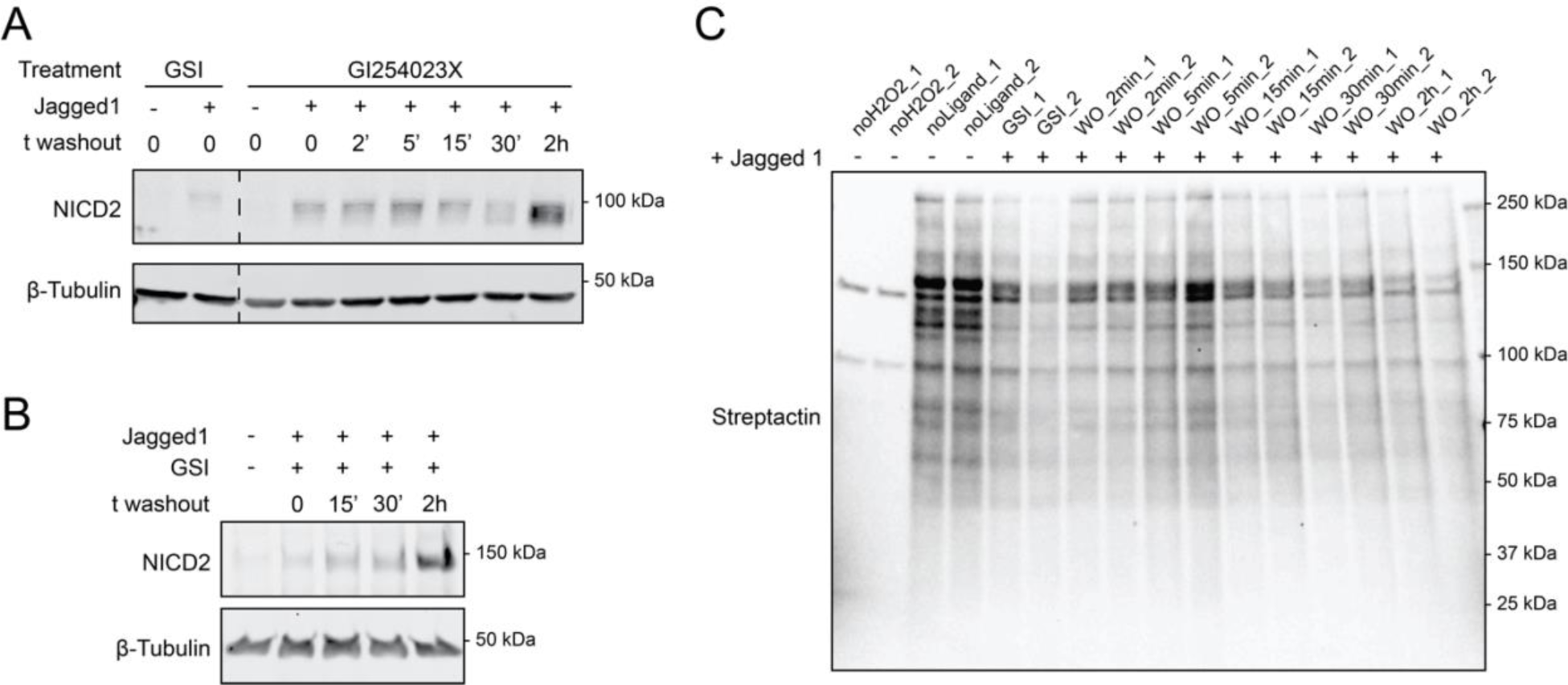
Notch2-APEX2 activation kinetics and biotinylation patterns, related to Fig. 1. (A) Western blot of Jag1-stimulated SVG-A cell lysates, probing for activated NICD2 as a function of time after removal of GI254023X (5 µM). (B) Western blot of Jag1-stimulated Notch2-APEX2 SVG-A cell lysates, probing for activated NICD2 as a function of time after removal of GSI (100 nM). (C) Streptactin blot of Jag1-stimulated SVG-A cell lysates, probing for proteins biotinylated by Notch2-APEX2 as a function of time after removal of GSI (WO).

**Fig. S3.**
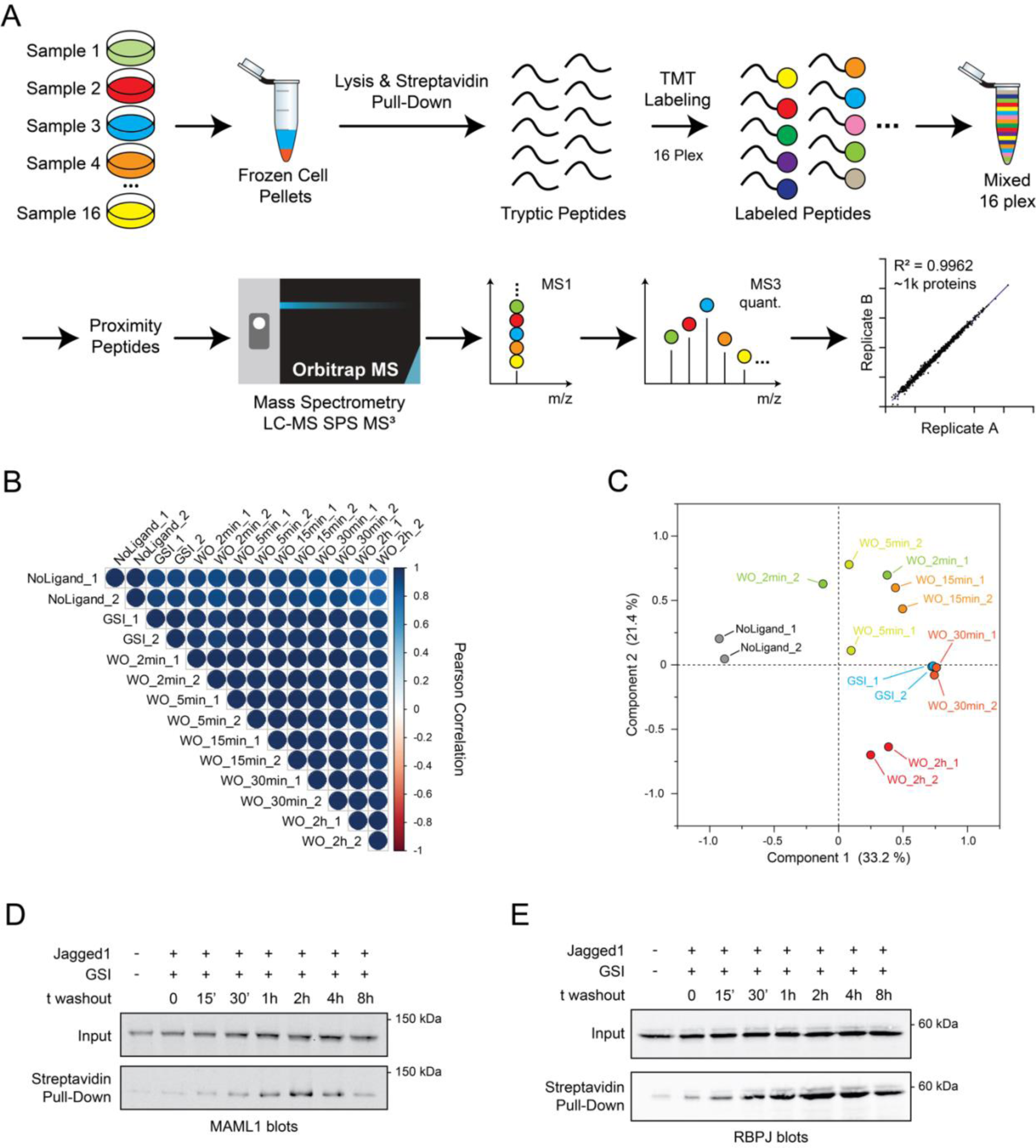
Notch2-APEX2 proximity labeling workflow and time-course reproducibility, related to Fig. 1. (A) Sample preparation and mass spectrometry workflow for analysis of the Notch2-APEX2 proximity labeling time course. After collection, samples were lysed and biotin-labeled proteins were purified on streptavidin beads in denaturing conditions. Recovered proteins were digested with trypsin, and then labeled using tandem mass tags (TMT, 16 plex) to enable quantitative mass spectrometric analysis of 16 different samples at the same time. (B) Pearson correlation matrix showing good reproducibility between internal replicates. (C) Principal Component Analysis (PCA) of the Notch2-APEX2 proximity labeling time-course. Each dot represents a sample and each color a time point. (D-E) Western blots of JAG1-stimulated Notch2-APEX2 knock-in SVG-A cells, probing for MAML1 (D) and RBPJ (E) as a function of time after GSI washout. Top: input; bottom: streptavidin pull-down after APEX2-mediated labeling.

**Fig. S4.**
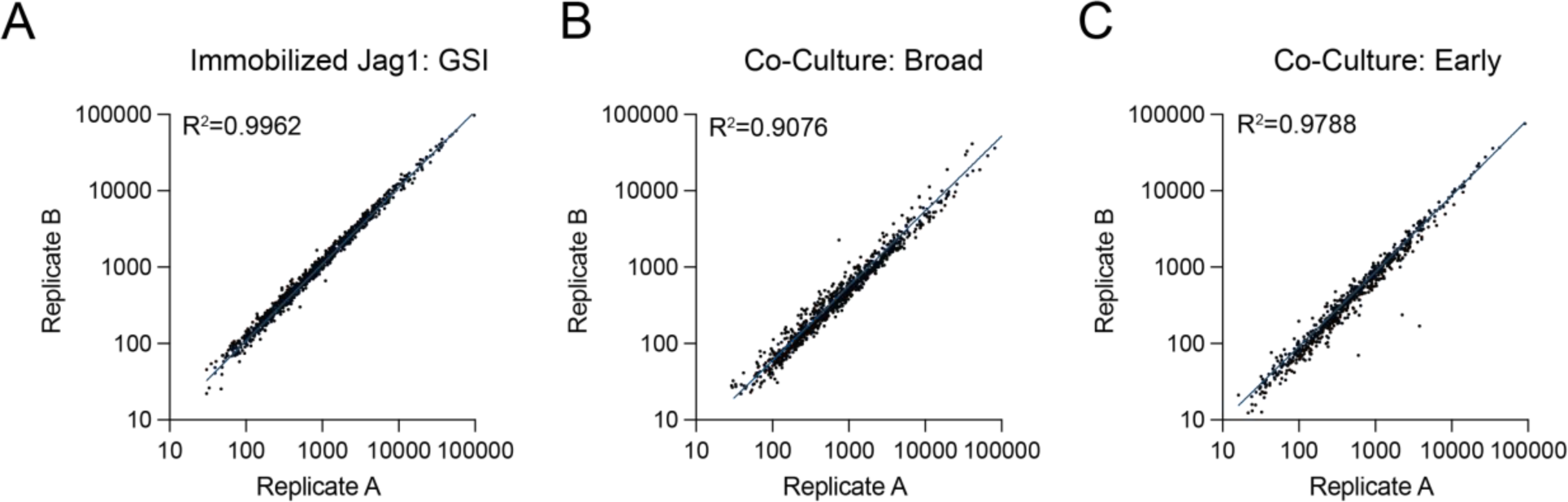
Replicate correlation analysis, related to Fig. 1. (A-C) Comparison of the correlation between replicates when Notch-APEX2 SVG-A cells are stimulated with immobilized Jag1 (A; panel duplicated from Fig. S3A) or with co-cultured Jag1-expressing A673 cells (B, C).

**Fig S5.**
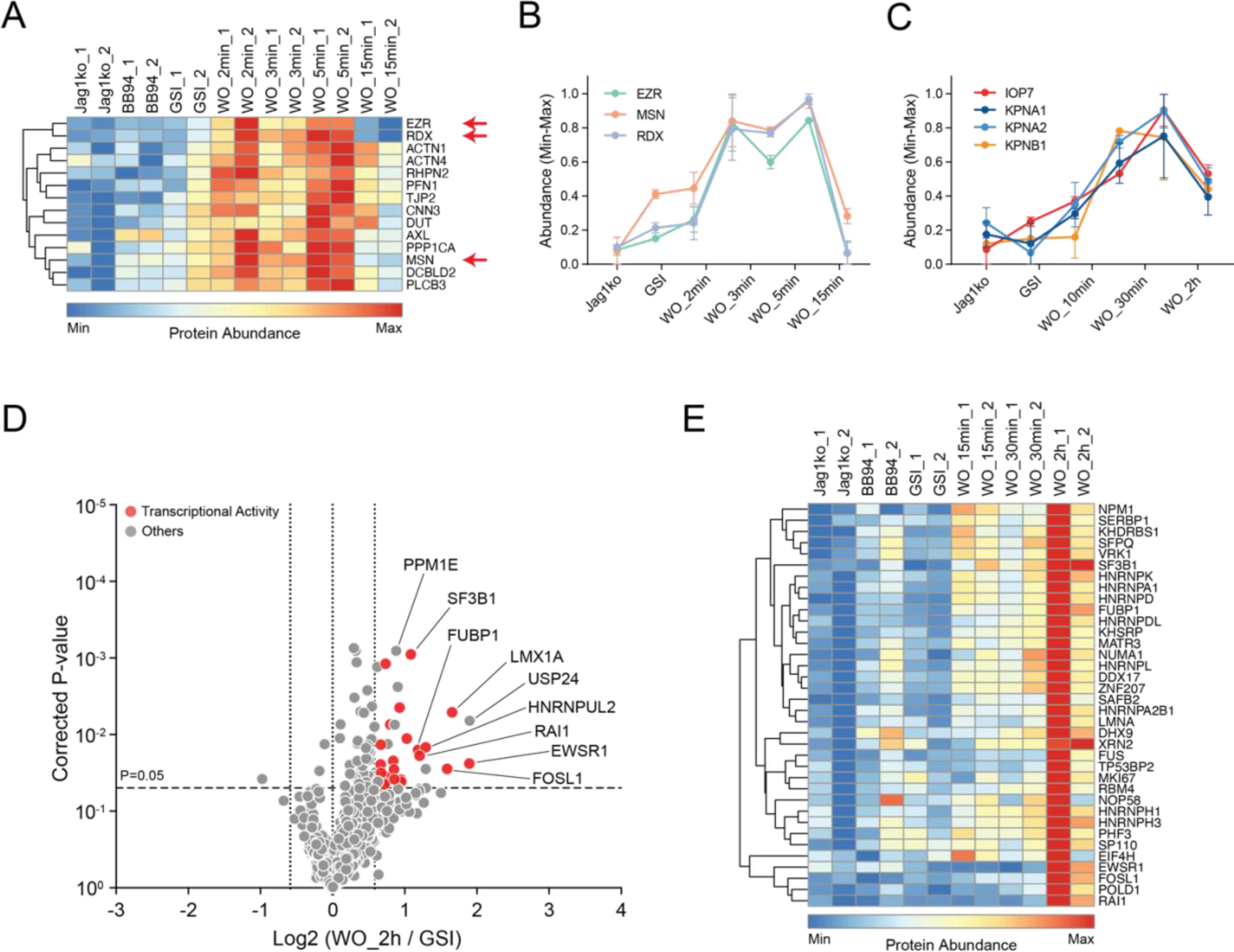
Proximity labeling results from co-culture of Notch2-APEX2 SVG-A cells with Jag1-expressing cells, related to Figs. 1-5. (A) Heatmap from the early time-course highlighting the ERM proteins Ezrin (EZR), Radixin (RDX), and Moesin (MSN) and proteins with similar labeling dynamics, in which labeling enrichment is observed in the 2-5 min time window. (B) Line plot showing labeling patterns of EZR, RDX, and MSN in the early time-course experiment. (C) Line plot showing labeling enrichment of the nuclear import proteins KPNA1, KPNA2, KPNB1, and IOP7, which reaches a maximum 30 min after washout. (D-E) Volcano plot (D) and heatmap (E) showing enrichment of nuclear proteins 2h after GSI washout in the broad time-course experiment.

**Fig. S6.**
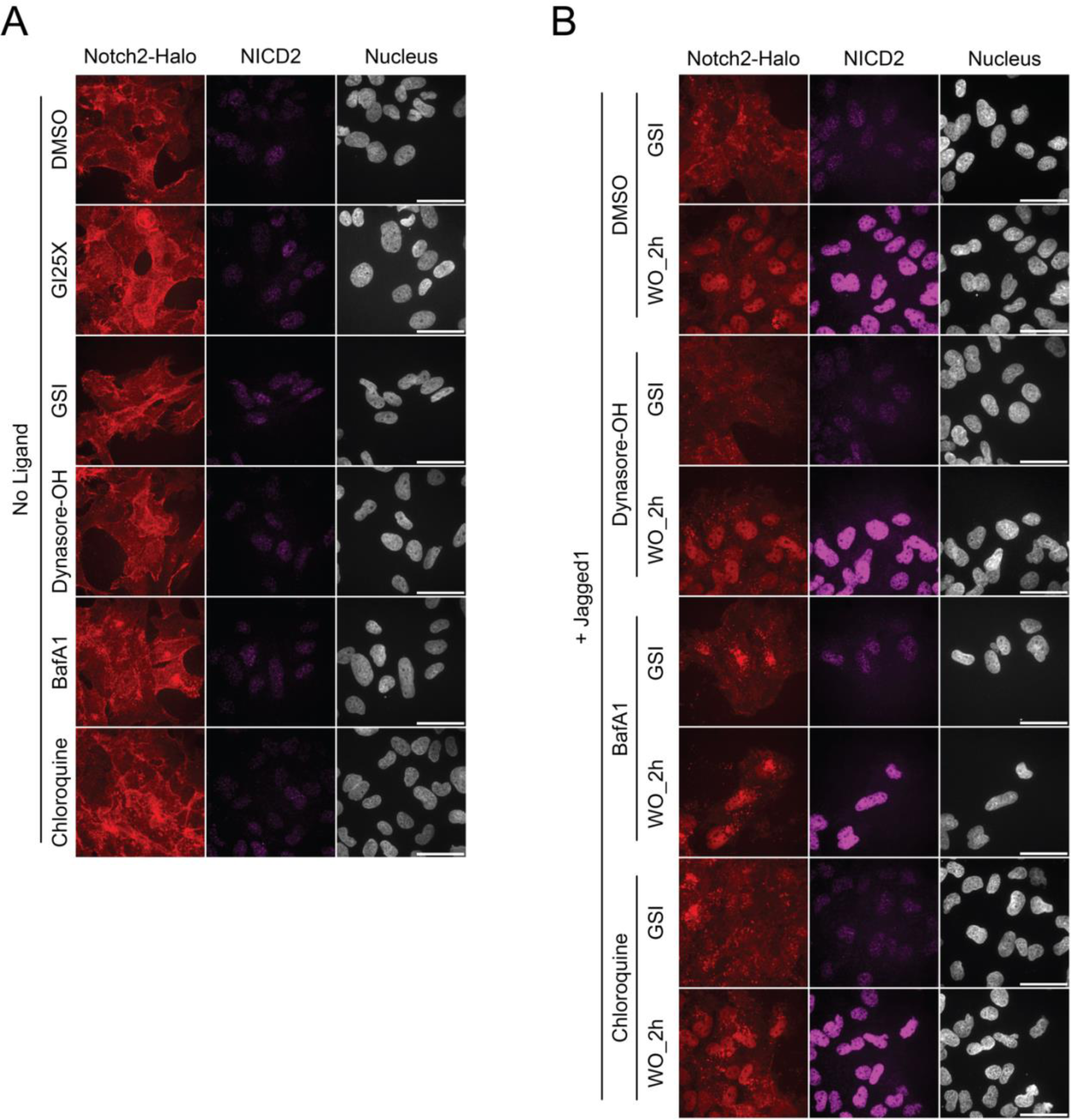
Effect of inhibition of endocytosis or vesicular acidification on NICD2 nuclear accumulation after GI254023X or GSI washout, related to Fig. 3. (A) Representative images of SVG-A Notch2-HaloTag cells showing the subcellular localization of Notch2-HaloTag (Notch2-Halo) and (S3-cleaved) NICD2 in the presence of DMSO, 5 µM GI254023X, 100 nM GSI, 20 µM hydroxy-dynasore, 25 nM bafilomycinA1 (BafA1), or 50 µM chloroquine in the absence of ligand stimulation. The HaloTag was labeled with JaneliaFluorX549 HaloTag ligand and NICD2 was stained with an anti-NICD2 primary antibody and anti-rabbit secondary antibody conjugated to Alexa Fluor 647. Nuclei were identified by DAPI staining. Scale bars: 20 µm. (B) Representative images of Jag1-stimulated SVG-A-Notch2-HaloTag cells showing the subcellular localization of Notch2-HaloTag (Notch2-Halo) and (S3-cleaved) NICD2 before and after removal of GSI (100 nM) in the absence or presence of hydroxy-dynasore (20 µM), bafilomycinA1 (BafA1; 25 nM), or chloroquine (50 µM). Nuclei were identified by DAPI staining. Scale bars: 20 µm.

**Fig. S7.**
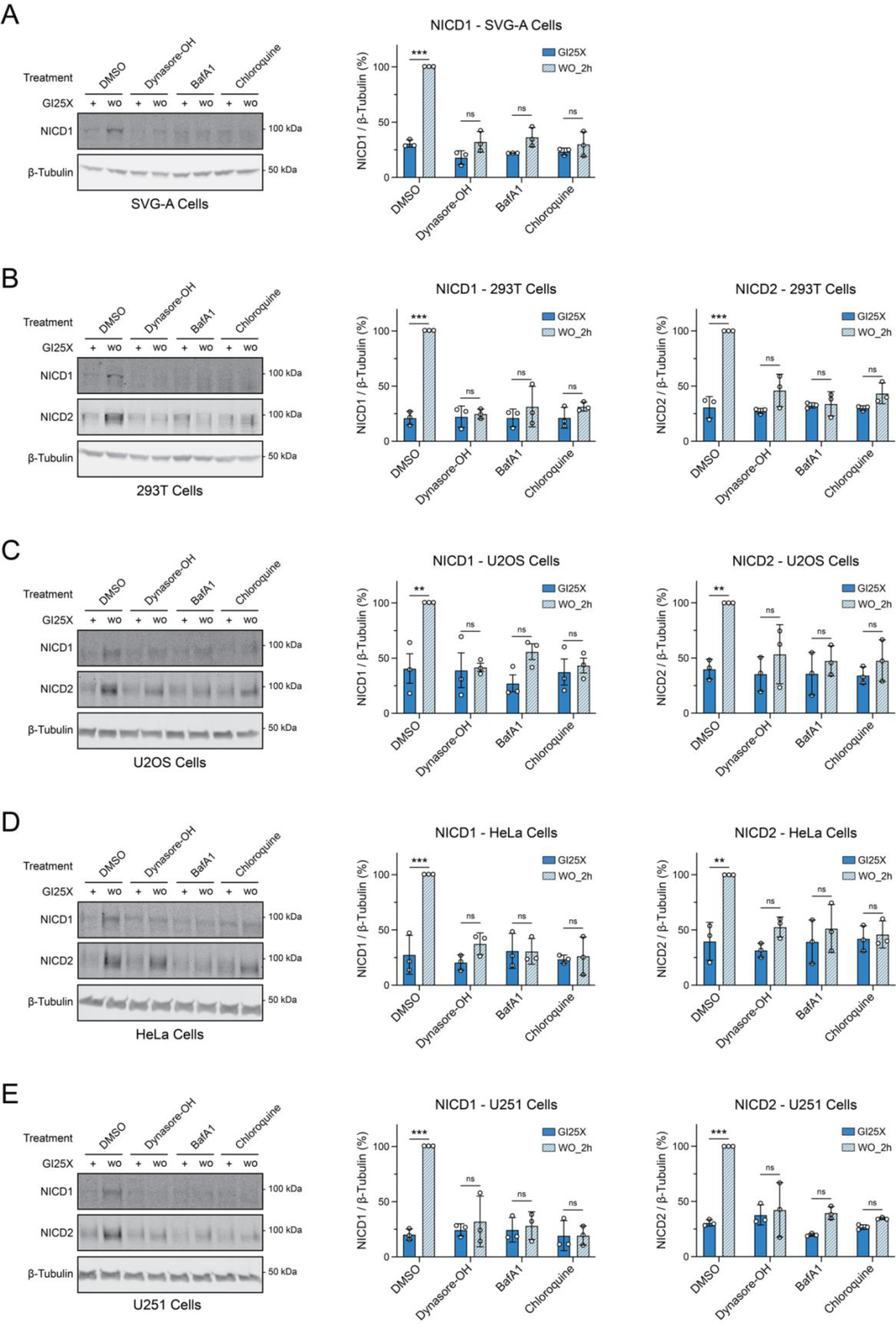
Effect of inhibition of endocytosis or vesicular acidification on NICD1 and NICD2 generation in different cell lines, related to Fig. 3. (A-E) Representative Western blot analysis (left) and quantifications (right) for NICD1 and/or NICD2 in Jag1-stimulated SVG-A (A), 293T (B), U2OS (C), HeLa (D), or U251 (E) cells 2 h after GI25X washout in the presence of different endocytosis and vesicular trafficking inhibitors. All data presented in this figure are from three independent experiments and are presented as mean ± SD. *p ≤ 0.05, *p ≤ 0.01 and ***p ≤ 0.001. Two-way ANOVA followed by Tukey’s pairwise comparison was used for statistical comparisons.

**Fig. S8.**
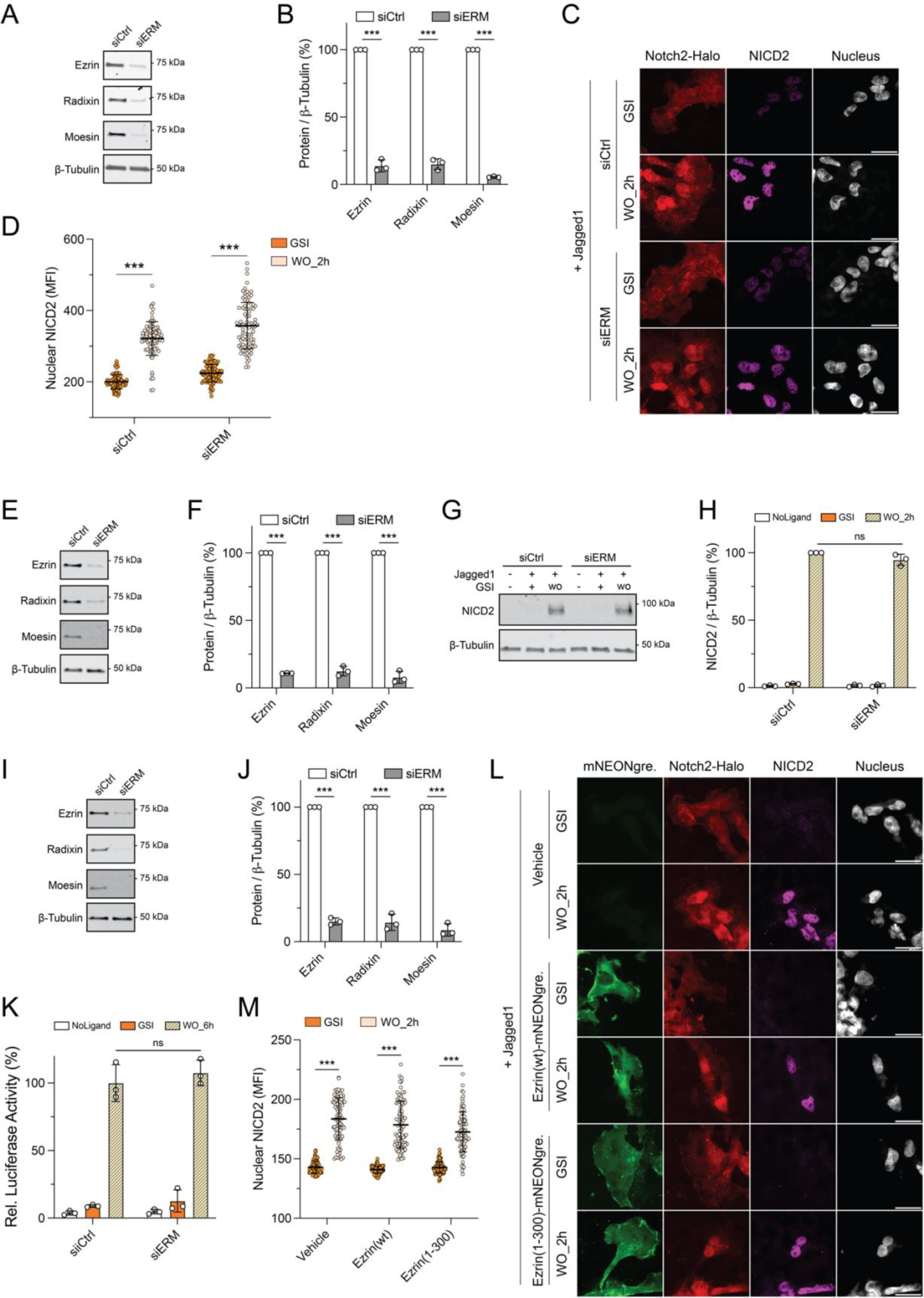
Effect of ERM inhibition on Notch2 signaling in SVG-A cells, related to Fig. 4. (A) Western blot and (B) quantification of band intensity to analyze siRNA-mediated knockdown of ERM proteins (n=3) for immunofluorescence experiments (C-D). (C) Representative images of Jag1-stimulated SVG-A Notch2-HaloTag cells showing the cellular localization of Notch2-HaloTag (Notch2-Halo) and NICD2 before and 2 h after removal of GSI (100 nM) as a function of siRNA knockdown of ERM (Ezrin, Radixin, Moesin) proteins. The HaloTag was labeled with JaneliaFluorX549 HaloTag ligand and NICD2 was stained with an anti-NICD2 primary antibody and anti-rabbit secondary antibody conjugated to Alexa Fluor 647. Nuclei were identified by DAPI staining. Scale bars: 20 µm. (D) Quantification of mean fluorescence signal intensity (MFI) in the nucleus for NICD2 from the imaging data in panel C (n=90 cells). (E) Western blot and (F) quantification of band intensity to analyze siRNA-mediated knockdown of ERM proteins (n=3) for experiments probing NICD2 production by Western blot (G-H). (G) Representative Western blot for NICD2 abundance before and after removal of GSI and (H) quantification of band intensity from the data in panel G (n=3). (I) Western blot and (J) quantification of band intensity to analyze siRNA-mediated knockdown of ERM proteins (n=3) for luciferase reporter assays shown in panel K. (K) Relative luciferase activity comparing the effect of siRNA-mediated knockdown of ERM proteins on Notch2 reporter gene activity in unstimulated cells, in Jag1-stimulated cells in the presence of GSI (100 nM), or 6 h after GSI removal (n=3). (L) Representative images showing the cellular localization of Notch2-Halotag (Notch2-Halo) and NICD2 before and after removal of GSI (100 nM) in Jag1-stimulated SVG-A Notch2-HaloTag cells transfected with vehicle, with wild-type Ezrin fused to mNeongreen, or with the Ezrin(1-300) dominant negative mutant fused to mNeongreen. Nuclei were identified by DAPI staining. Scale bars: 20 µm. (M) Quantification of mean fluorescence signal intensity (MFI) in the nucleus for NICD2 using the imaging data of panel L (90 cells). Only cells positive for expression of either wild-type or dominant negative mutant of Ezrin were selected for quantification. All data presented in this figure are from three independent experiments (n=3) and are presented as mean ± SD. *p ≤ 0.05, *p ≤ 0.01 and ***p ≤ 0.001. Two-way ANOVA followed by Tukey’s pairwise comparison was used for statistical comparisons.

**Fig. S9.**
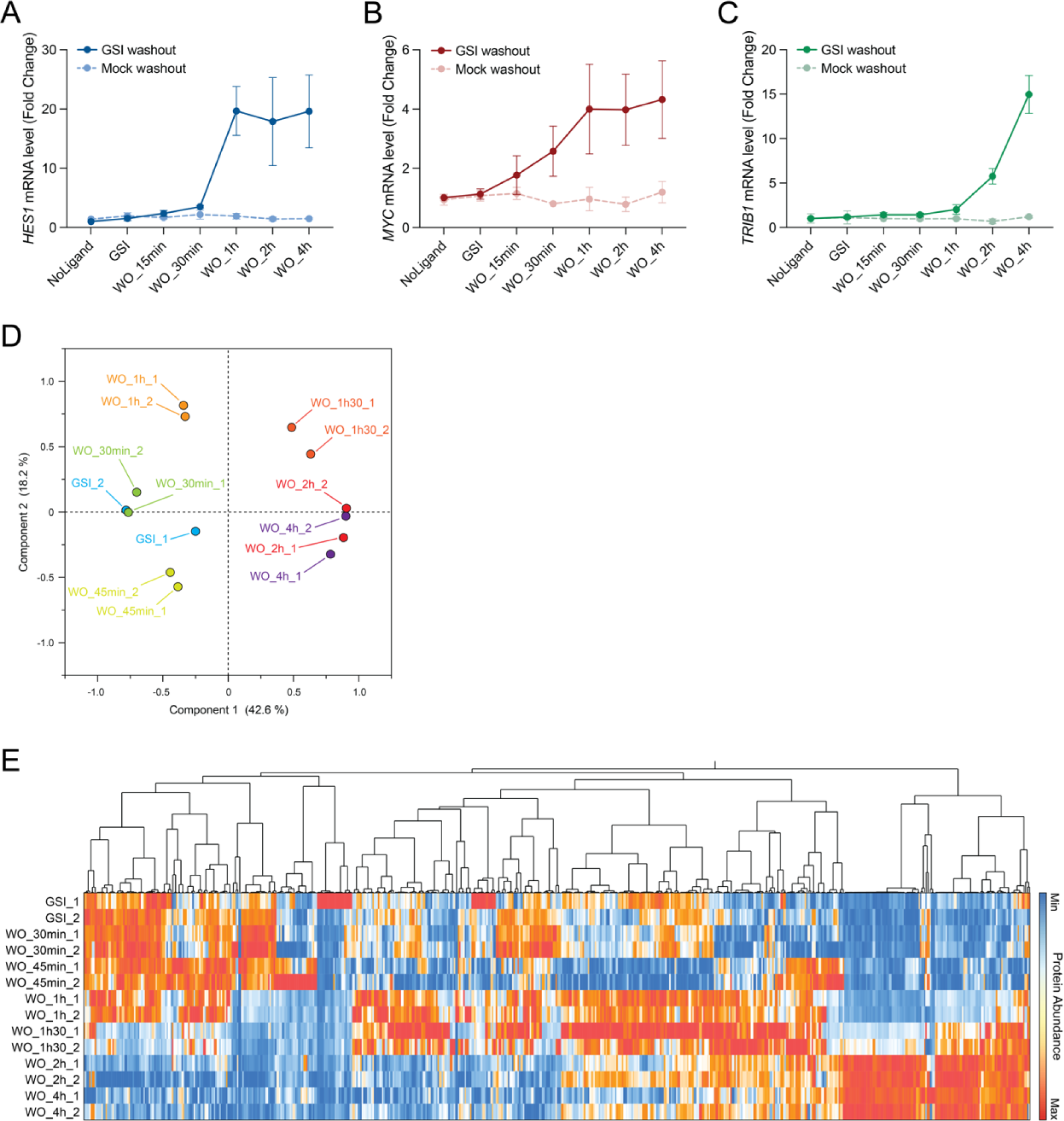
Notch2-APEX2 time-resolved proximity labeling focused on nuclear entry, related to Fig. 5. (A-C) qRT-PCR for the Notch target genes *HES1* (A), *MYC* (B), and *TRIB1* (C) in parental SVG-A cells as a function of time after GSI washout. Data presented in this panel set are from three independent experiments and are presented as mean ± SD. (D) Principal Component Analysis (PCA) of enrichment profiles of the proteins identified in the Notch2-APEX2 proximity labeling time-course centered around NICD2 nuclear entry. Each dot represents a sample and each color a time point. (E) Heatmap of hierarchical clustering of Notch2-APEX2 proximity labeling as a function of time after GSI washout using plated Jag1 with timepoints centered around NICD2 nuclear entry. Clustering of the relative abundance of each identified protein (columns) as a function of time (rows) was performed based on Ward’s minimum variance method. The color scheme representing the relative abundance for each protein (minimum to maximum) is shown on the right.

**Data File S1. List of proteins identified and their enrichment levels for Notch2-APEX2 using immobilized ligand with GSI washout.**

**Data File S2. List of proteins identified and their enrichment levels for Notch2-APEX2 using co-culture with A673 cells - Broad.**

**Data File S3. List of proteins identified and their enrichment levels for Notch2-APEX2 using co-culture with A673 cells - Early.**

**Data File S4. List of proteins identified and their enrichment levels for Notch2-APEX2 using immobilized ligand with GSI nuclear centered washout.**

**Table S1.** List of key reagents used in this study.

| **Reagent type** | **Designation** | **Source of reference** | **Identifiers** | **Additional information** |
| --- | --- | --- | --- | --- |
| Antibody | Mouse monoclonal anti β-Tubulin (clone D3U1W) | Cell Signaling Technology | RRID: AB_2715541 / Catalog: #86298S | Western blot (1:2000) |
| Antibody | Rabbit monoclonal anti GAPDH (clone 14C10) | Cell Signaling Technology | RRID: AB_561053 / Catalog: #2118S | Western blot (1:2000) |
| Antibody | Rabbit monoclonal anti Notch2 (clone D76A6) | Cell Signaling Technology | RRID: AB_10693319 / Catalog: #5732S | Western blot (1:1000) |
| Antibody | Rabbit anti NICD2 | Eli Lilly | See (41) | Western blot (1:1000) & Immunofluorescence (1:2500) |
| Antibody | Rabbit monoclonal anti NICD1 (clone D3B8) | Cell Signaling Technology | RRID: AB_2153348 / Catalog: #4147S | Western blot (1:1000) |
| Antibody | Mouse monoclonal anti HA-tag (clone 6E2) | Cell Signaling Technology | RRID: AB_10691311 / Catalog: #2367S | Western blot (1:500) |
| Antibody | Rabbit monoclonal anti RBPJ (clone D10A4) | Cell Signaling Technology | RRID: AB_2665555 / Catalog: #5313S | Western blot (1:1000) |
| Antibody | Mouse monoclonal anti mNeonGreen (clone 6G6) | Chromotek | RRID: AB_2827566 / Catalog: #32F6 | Western blot (1:500) |
| Antibody | APC anti Notch2 | BioLegend | RRID: AB_10662412 / Catalog: #348306 | Flow Cytometry (2 μL for 500 000 cells) |
| Antibody | APC Mouse Isotype | BioLegend | RRID: AB_2891178 / Catalog: #400221 | Flow Cytometry (2 μL for 500 000 cells) |
| Antibody | Alexa Fluor Plus 647-conjugated donkey anti-rabbit polyclonal secondary | ThermoFisher Scientific | RRID: AB_2762835 / Catalog: #A32795 | Immunofluorescence (1:1000) |
| Cell line | SVG-A | Walter J. Atwood, Brown University | RRID: CVCL_5G13 |  |
| Cell line | Notch2 knockout SVG-A cell line | This study |  |  |
| Cell line | Notch2-APEX2 knockin SVG-A cell line | This study |  |  |
| Cell line | 293T | ATCC | RRID: CVCL_0063 / Catalog: #CRL-3216 |  |
| Cell line | U2OS | ATCC | RRID: CVCL_0042 / Catalog: #HTB-96 |  |
| Cell line | HeLa | ATCC | RRID: CVCL_0030 / Catalog: #CCL-2 |  |
| Cell line | U251 | Sigma | RRID: CVCL_0021 / Catalog: #1610381 |  |
| Reagent | Biotin-Phenol | Iris Biotech | Catalog: #LS-3500 |  |
| Reagent | Hydrogen Peroxide | Sigma | Catalog: #H1009-5ML |  |
| Reagent | Pierce Streptavidin magnetic beads | ThermoFisher Scientific | Catalog: #88817 |  |
| Reagent | Streptactin-HRP | Bio-rad | Catalog: #1610381 | Western blot (1:1000) |
| Reagent | JaneliaFluorX549 HaloTag ligand | Luke Lavis, Janelia Research Campus |  | HaloTag labeling (100 nM) |
| siRNA | siCtrl | ThermoFisher Scientific | AM4611 |  |
| siRNA | siEZR (Ezrin) | ThermoFisher Scientific | s14795 |  |
| siRNA | siRDX (Radixin) | ThermoFisher Scientific | s11899 |  |
| siRNA | siMSN (Meosin) | ThermoFisher Scientific | s8984 |  |

**Table S2:**
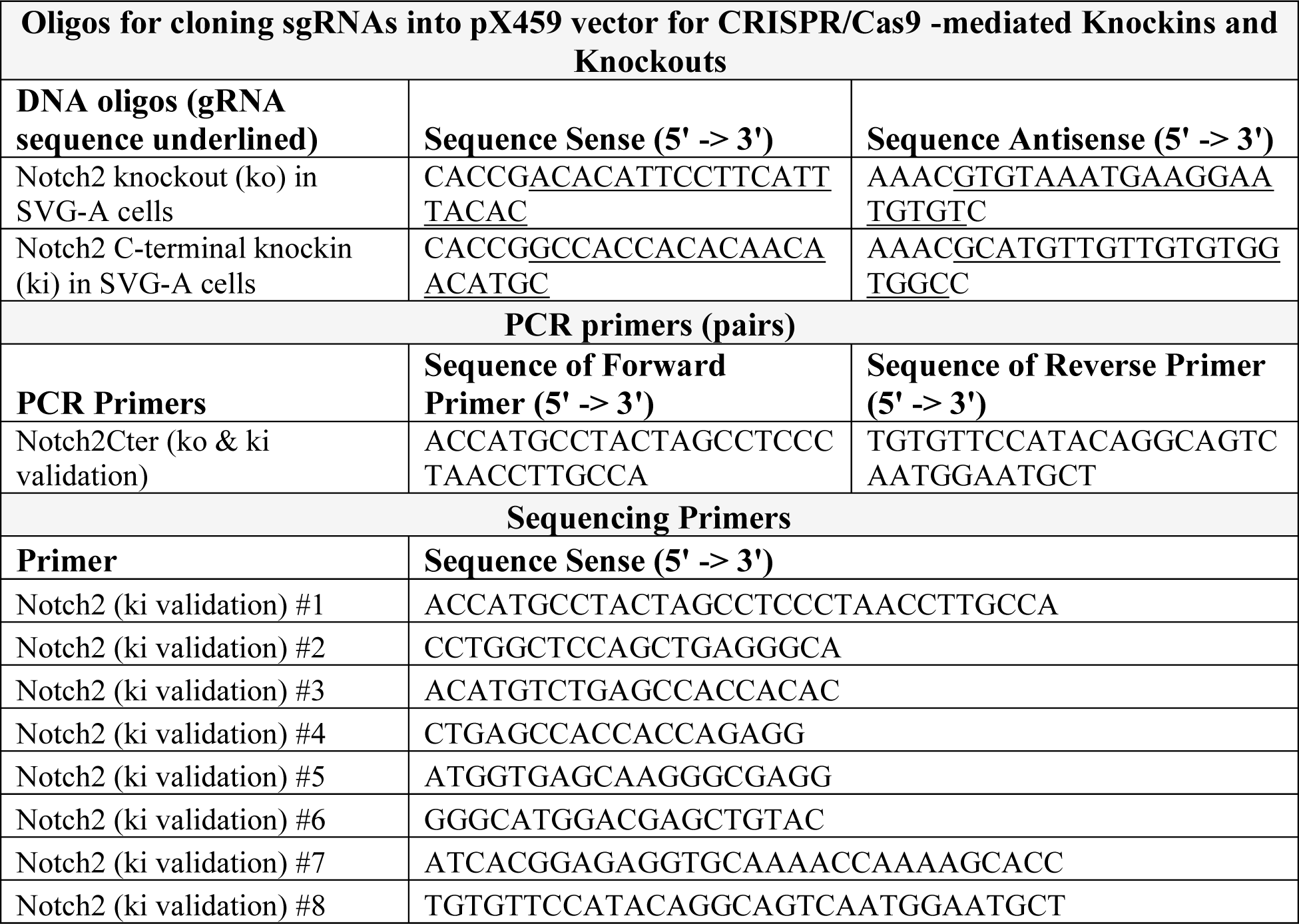
List of CRISPR reagents used in this study.

**Table S3.**
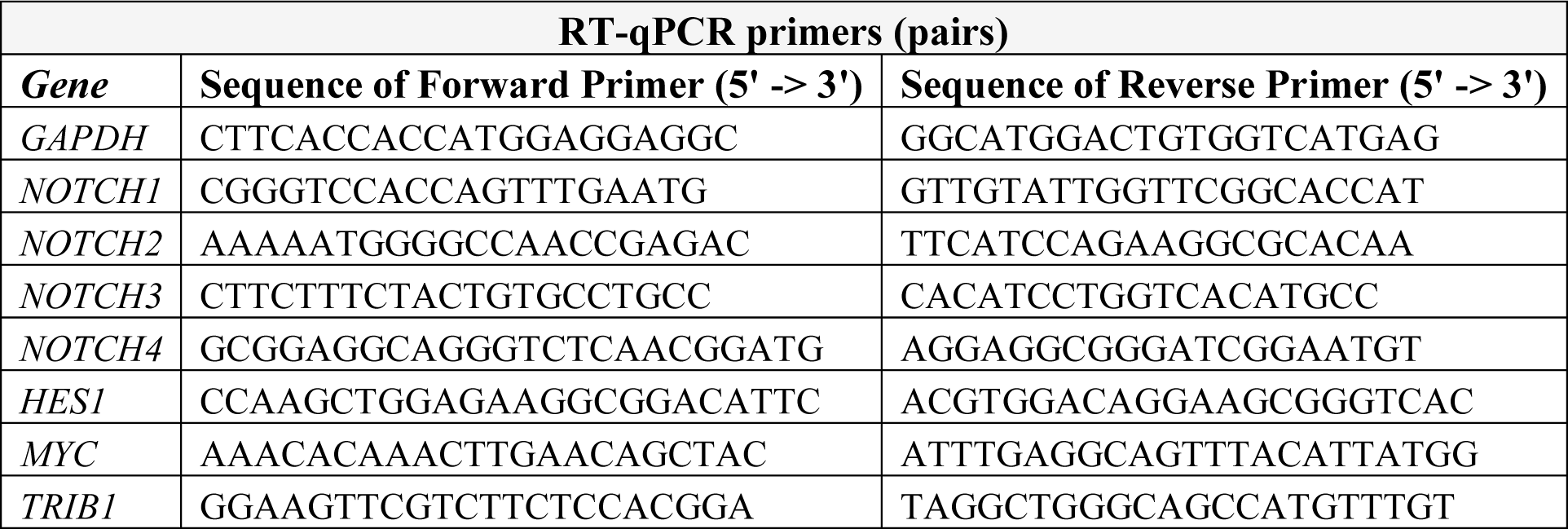
List and sequence of qRT-PCR primers used in this study.

